# Previously unmeasured genetic diversity explains part of Lewontin’s paradox in a *k*-mer-based meta-analysis of 112 plant species

**DOI:** 10.1101/2024.05.17.594778

**Authors:** Miles Roberts, Emily B. Josephs

## Abstract

At the molecular level, most evolution is expected to be neutral. A key prediction of this expectation is that the level of genetic diversity in a population should scale with population size. However, as was noted by Richard Lewontin in 1974 and reaffirmed by later studies, the slope of the population size-diversity relationship in nature is much weaker than expected under neutral theory. We hypothesize that one contributor to this paradox is that current methods relying on single nucleotide polymorphisms (SNPs) called from aligning short reads to a reference genome underestimate levels of genetic diversity in many species. To test this idea, we calculated nucleotide diversity (*π*) and *k*-mer-based metrics of genetic diversity across 112 plant species, amounting to over 205 terabases of DNA sequencing data from 27,488 individual plants. We then compared how these different metrics correlated with proxies of population size that account for both range size and population density variation across species. We found that our population size proxies scaled anywhere from about 3 to over 20 times faster with *k*-mer diversity than nucleotide diversity after adjusting for evolutionary history, mating system, life cycle habit, cultivation status, and invasiveness. The relationship between *k*-mer diversity and population size proxies also remains significant after correcting for genome size, whereas the analogous relationship for nucleotide diversity does not. These results suggest that variation not captured by common SNP-based analyses explains part of Lewontin’s paradox in plants.

**Lay Summary:** Even after many revolutions in our ability to sequence and understand DNA, many important biological questions remain unsolved. One such problem is Lewontin’s paradox, named after Richard Lewontin who first described it in 1974. The core of the paradox is a simple idea: species with more individuals should be more genetically diverse. The reasoning is that more individuals means more replication of DNA, and thus more opportunities for mutation to create new variation. However, species that differ massively in population size often have similar diversity levels. Lewontin’s paradox has several potential, previously investigated mechanisms but what if one contributor is simply that our measurements of genetic diversity are off? Most studies estimate diversity by comparing sample genomes to a standard reference genome. While this approach is useful, it is impossible to measure variation in DNA that is not represented in the reference - a phenomenon known as reference bias. We estimate metrics of diversity that are free of reference-bias and re-investigate Lewontin’s paradox in plants. Overall, we find that reference-free diversity metrics scale more with population size, compared to the reference-biased approach. While it is unlikely that reference-bias fully explains Lewontin’s paradox, our analyses suggest that reference-bias plays an important role.

## Introduction

Understanding the determinants of genetic diversity within populations is key to informing species conservation [Cole, 2003] and breeding efforts [Sanchez et al., 2023]. However, most species have far less genetic diversity (commonly estimated as pairwise nucleotide diversity, *π*) than expected [Frankham, 2012, Corbett-Detig et al., 2015, Buffalo, 2021]. If we assume that the vast majority of genetic variants are neutral, then the determinants of genetic diversity are encapsulated in neutral theory [Kimura, 1983]: *E*[*π*] *≈* 4*N_e_µ*, where *E*[*π*] is the expected level of genetic diversity, *N_e_* is the effective size of a population, and *µ* is the mutation rate per base pair per generation. Mutations rates vary relatively little across species (Cagan et al. [2022], Bergeron et al. [2023], reviewed in Quiroz et al. [2023]), while the total number of individuals in a species varies massively [Buffalo, 2021]. Thus, under neutral theory, population size should be a strong determinant of genetic diversity and species with larger population sizes should be more diverse. However, even some of the most abundant species studied to date have low genetic diversity compared to neutral theory expectations. For example, *Drosophila simulans* has an estimated population size *>* 10^14^ and a diversity of *π ≈* 0.01, but an expected diversity of *π >* 0.1 [Buffalo, 2021]. This mismatch between expected and observed levels of neutral diversity across populations of varying size is known as Lewontin’s paradox, named after Richard Lewontin who first described the phenomenon [Lewontin, 1974].

The potential mechanisms underlying Lewontin’s paradox have been reviewed extensively [Leffler et al., 2012, Slotte, 2014, Ellegren and Galtier, 2016, Charlesworth and Jensen, 2022]. Multiple selective and demographic processes likely contribute to Lewontin’s paradox; however, determining the relative importance of these processes remains a contentious area of research. The two most explored mechanisms are historic population size changes [Charlesworth and Jensen, 2022] and linked selection - whereby fixation or purging of selected alleles causes the loss of linked neutral alleles [Kojima and Schaffer, 1964, 1967, Smith and Haigh, 1974, Charlesworth et al., 1993, Charlesworth, 1994]. Linked selection is expected to reduce diversity more in regions of lower recombination and higher functional density [Slotte, 2014]. Thus, many studies have focused on measuring the correlations of recombination rate or functional density with either intraspecific or interspecific diversity, often observing significant correlations [Tenaillon et al., 2001, Hellmann et al., 2003, Nordborg et al., 2005, Roselius et al., 2005, Branca et al., 2011, Paape et al., 2012, Corbett-Detig et al., 2015, Silva-Junior and Grattapaglia, 2015, Wang et al., 2016, Phung et al., 2016, Mackintosh et al., 2019]. However, not all studies observe strong correlations between recombination and diversity [Schmid et al., 2005, Roselius et al., 2005, Flowers et al., 2012, Wang et al., 2016], especially studies focused on plant species (reviewed in Slotte [2014]), and such correlations could be explained by an association between recombination and mutation [Hellmann et al., 2003] (though the evidence for this is mixed, see Mackintosh et al. [2019]). There is also both empirical and theoretical evidence that linked selection is unlikely to explain the entirety of Lewontin’s paradox, suggesting that demographic factors play an important role [Coop, 2016, Buffalo, 2021, Charlesworth and Jensen, 2022].

There are three main types of demographic changes proposed to contribute to Lewontin’s paradox: contractions, expansions, and cyclical population size changes [Charlesworth and Jensen, 2022]. Population contractions cause loss of diversity. Thus, if many species’ populations recently contracted (due to human activity, for example), then their contemporary diversity would be much lower than expected from their pre-contraction population sizes [Exposito-Alonso et al., 2022]. Recent population expansions could cause a similar mismatch. Because it takes many generations for populations to accumulate diversity compared to the timescale of typical expansions, contemporary diversity levels for an expanded population would be much smaller than expected from a post-expansion population size [Peart et al., 2020, Charlesworth and Jensen, 2022]. For a similar reason, species that have seasonal variation in their population sizes will also tend to have diversity levels closer to what one would expect based on their minimum size rather than their peak size [Wright, 1940]. Studies investigating Lewontin’s paradox would ideally try to jointly infer these demographic histories alongside selective factors in natural populations. However, issues of model complexity and identifiability often prevent such joint estimation [Johri et al., 2020, 2022b,a], suggesting further explorations of Lewontin’s paradox will require new approaches.

One potential, but rarely explored, contributor to Lewontin’s paradox is that current methods for estimating genetic diversity systematically underestimate the true levels of genetic diversity in most populations. Lewontin’s original observations and earlier studies on the population size-diversity relationship were based on allozymes, which detect variants in protein sequences [Lewontin, 1974, Nei and Graur, 1984]. More recent studies measure diversity using SNPs at more neutral four-fold degenerate sites (i.e. sites where mutations do not affect protein sequences) in DNA and generally observe greater within-species diversity and between-species divergence compared to allozymes [Li and Sadler, 1991, Makalowski and Boguski, 1998, Bazin et al., 2006, Piganeau and Eyre-Walker, 2009]. However, current SNP-based methods are not perfect either and there is significant evidence that SNPs capture a biased and incomplete picture of genetic diversity. First, calling SNPs typically requires aligning reads to a reference genome, meaning any SNPs in regions that are not present or highly diverged from the reference genome will be excluded from analysis and thus downwardly bias diversity estimates [Golicz et al., 2020, Buffalo, 2021]. This downward bias is typically assumed to have little effect on the qualitative relationship between diversity and *N_e_* [Buffalo, 2021], but recent pangenomic studies have uncovered troves of non-reference variation across a variety of species (Ebler et al. [2022], Rice et al. [2023], reviewed in Bayer et al. [2020]). Second, many other classes of genetic variants contribute to genetic diversity besides SNPs, and SNPs can actually be a cryptic sign of larger-scale variation. For example, a large fraction of heterozygous SNP calls in *Arabidopsis thaliana* are actually the result of structural variation [Jaegle et al., 2023]. Finally, previous meta-analyses of population size and diversity data rely on scraping diversity estimates from previously published studies (Frankham [2012], Buffalo [2021], except see Corbett-Detig et al. [2015]). However, many studies report inaccurate SNP calls and diversity estimates due to errors in the handling of missing data [Korunes and Samuk, 2021, Schmidt et al., 2021, Sopniewski and Catullo, 2024] and may filter genotype calls differently, making comparisons across species difficult. Overall, errors in diversity calculations and omission of diversity in genomic regions that are either difficult or impossible to align to could partially explain Lewontin’s paradox. Re-analyzing whole genome sequencing data with a common pipeline would make diversity estimates across species more comparable and easier to interpret [Buffalo, 2021, Mirchandani et al., 2024].

One useful pangenomics tool for measuring non-reference variation that is readily applicable to common short-read datasets is the *k*-mer. *k*-mers are subsequences of length *k* derived from a larger sequence and they have a long history of use in computer science [Shannon, 1948], genome assembly [Turner et al., 2018], metagenomics [Benoit et al., 2016], and quantitative genetics [Rahman et al., 2018, Voichek and Weigel, 2020, Kim et al., 2020, Mehrab et al., 2021]. Recent studies have also demonstrated the utility of *k*-mers for measuring heterozygosity and genetic differences between individuals (commonly referred to as “dissimilarity” measures, Ondov et al. [2016], Vurture et al. [2017], Ranallo-Benavidez et al. [2020], VanWallendael and Alvarez [2022]). Typical analysis of *k*-mers involves only counting the presence/absence and/or frequencies of all *k*-mers in a set of reads, without aligning the reads to any reference, then deriving measures of genetic difference from such counts [Benoit et al., 2016]. Avoiding alignment allows one to incorporate sequences that would otherwise be omitted for lack of alignment to a reference genome.

We revisited Lewontin’s paradox in plants using *k*-mer-based measures of genetic difference, aiming to test whether the inclusion of non-reference variation could partially resolve Lewontin’s paradox. We compared how *k*-mer dissimilarity and typical SNP-based estimates of nucleotide diversity correlated with population size proxies across a large panel of plant species - all processed through the same bioinformatic pipeline. Our expectation was that if *k*-mers are better at capturing genomic variation than SNPs, *k*-mer dissimilarity would scale more rapidly with population size compared to nucleotide diversity.

## Materials and Methods

Our entire analysis is packaged as a snakemake workflow stored here: https://github.com/milesroberts-123/tajimasDacrossSpecies. This workflow includes the code to reproduce all of the steps individually explained below, along with instructions on how to run the code, and yaml files describing the exact configurations of software we used at each step. It also includes an example directed acyclic graph showing the order of steps a typical sample is processed through. The code detailing all initial, exploratory, and confirmatory data analyses as well as figure creation can be found as an R-markdown file in the github repository. The parameters for each software were kept constant across all datasets (except occasionally for the “–ploidy” parameter in GATK HaplotypeCaller) to ensure that variation in bioinformatic processing did not bias our results. All statistical analyses used R v4.2.2 [R Core Team, 2022] and all color palettes used in figure creation come from the scico R package [Pedersen and Crameri, 2023] to ensure color-blind accessibility.

### Population-level sequencing data collection

We started by building a list of species with high quality, publicly available reference genomes as well as population-level sequencing data. The source for the genome assembly and annotation used for each species in this study is listed in Table S1. We first downloaded all genomes in Phytozome (https://phytozome-next.jgi.doe.gov/) with unrestricted data usage. We then downloaded all genomes for species from Ensembl plants (https://plants.ensembl.org/index.html) that were not already represented in Phytozome. Next, we downloaded genomes for additional species from the NCBI genome database (https://www.ncbi.nlm.nih.gov/genome/) that were not already present in either Phytozome or Ensembl and met all of the following criteria:

- matched filters: eukaryotic, plants, land plants, and exclude partial
- included assemblies of nuclear DNA (i.e. not just plastid genomes)
- included annotations of coding sequences

We also downloaded a genome for *Nicotiana tabaccum* from the Sol genomics network (https://solgenomics.net/). Finally, we omitted any species that had at least one chromosome longer than 2^29^ bp (about 512 Mb) from all downstream analyses because tabix indexing, which is often utilized for SNP-calling pipelines, does not support chromosomes exceeding this length. In the end, we were left with genome assemblies and annotations for 112 plant species (see Table S1).

Note that, similar to previous studies [Corbett-Detig et al., 2015, Buffalo, 2021], many of the plant species in this set of 112 are domesticated (see Table S1). This means that many of the species in our dataset have likely undergone recent demographic changes. However, we do not expect this to contribute to differences in the relationships between population size and nucleotide or *k*-mer diversity, because past demography affects both nucleotide and *k*-mer diversity. Furthermore, we include cultivation status in our downstream modeling to help account for systematic differences between cultivated and wild species (see **Statistical analysis**).

For each species with a reference genome, we searched for DNA-seq runs in the National Center for Biotechnology Information’s Sequence Read Archive (SRA) with a name in the organism field that matched the species name (e.g. search for Arabidopsis lyrata[Organism] to get *Arabidopsis lyrata* runs). We downloaded the run info for each search and found the study with most sequenced individuals for inclusion in our analysis. Most datasets came from individual studies, with the exception of *Zea mays*, which included several studies described in Bukowski et al. [2017]. The datasets used for each species are listed in Table S1.

We limited the size of each species’ dataset to no more than 7.5 *×* 10^12^ bp and no more than 1200 individuals because this defined the amount of data our workflow could process without the peak memory limit exceeding 50 TB and the time limit for genotype calling exceeding 7 days. If a species’ dataset exceeded either 1200 individuals or 7.5 *×* 10^12^ bp, we randomly downsampled runs such that both of these limits were satisfied.

We downloaded the SRA runs associated with each individual using the SRA toolkit (v2.10.7), then trimmed low-quality base calls with fastp (v0.23.1, Chen et al. [2018]), requiring a minimum quality score of 20 and a minimum read length of 30 base pairs. For each species, we summarized the results of fastp trimming using multiqc (v1.18, Ewels et al. [2016]). After trimming, any fastq files that were technical replicates of the same individual were concatenated. Concatenated fastq files were then processed through two different workflows: SNP-calling and *k*-mer counting.

### Single-nucleotide polymorphism calling

We aligned sequencing reads for each individual to their respective reference genome using BWA MEM (v0.7.17, Li and Durbin [2009], Li [2013]), sorted the resulting BAM files with samtools (v1.11, Danecek et al. [2021]), and marked optical duplicates with picardtools (picard-slim v2.22.1, Institute [2019]). Next, we called SNPs with GATK HaplotypeCaller (v4.1.4.1, McKenna et al. [2010], Poplin et al. [2018]). We varied the –ploidy parameter for HaplotypeCaller between species depending on the actual ploidy recorded in the literature and whether individual subgenome assemblies were available. However, the vast majority of species in our dataset had a –ploidy paramter of 2. We restricted genotype calling to only 4-fold degenerate sites within the nuclear genome, as identified by degenotate (v1.1.3, Mirchandani et al. [2024]), to focus solely on neutral diversity. Runs for each species were then combined with GATK GenomicsDBImport, then genotyped with GATK GenotypeGVCFs, including invariant sites as done in Korunes and Samuk [2021]. Variant and invariant sites were separated with bcftools (v1.17, Danecek et al. [2021]) and then filtered separately, as recommended by Korunes and Samuk [2021]. Variant sites were removed from our analyses if they met at least one of the following criteria: number of alleles *>* 2, indel status = TRUE, fraction of missing genotypes *>* 0.2, QD *<* 2.0, QUAL *<* 30.0, MQ *<* 40.00, FS *>* 60.0, HaplotypeScore *>* 13.0, MQRankSum *<* −12.5, and ReadPosRankSum *<* −8.0 [Caetano-Anolles, 2023]. For each species, we also required that each variant site have a minimum read depth of 5, but no more than 3 times the genome-wide average read depth at variant sites for that species. Meanwhile, invariant sites were removed from our analyses if they met at least one of the following criteria: QUAL *>* 100.0, read depth *≤* 5, or read depth *≥* 3 times the genome-wide average read depth at invariant sites for that species. Finally, invariant and variant sites were concatenated into a single VCF file per scaffold using bcftools. For *Brassica napus* and *Miscanthus sinensis*, scaffolds named “LK032656” (195,249 bp) and “scaffold04645” (2,838 bp), respectively, were omitted from our analyses because an error in SLURM job cancellation caused snakemake to prematurely delete intermediate files for these scaffolds. It is worth noting that different choices of genotype callers and filtering parameters could lead to different estimates of nucleotide diversity. However, our workflow is representative of SNP calling workflows used in many published population genetic analyses.

Using the SNP genotypes called from our pipeline, we then calculated genome-wide average nucleotide diversity at four-fold degenerate sites (*π̅*) using the filtered set of variant and invariant sites. To do this, we first calculated heterozygosity at each four-fold degenerate site (*i*) according to Hahn [2018]:

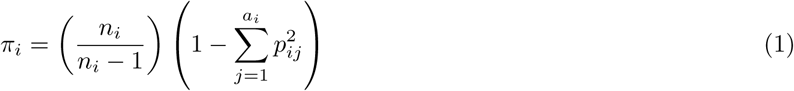

where *n_i_* is the number of sequenced chromosomes with non-missing genotypes for site *i*, *a_i_* is the number of alleles for site *i*, and *p_ij_* is the frequency of the *j*th allele at site *i*. For each invariant site, the equation reduces to *π_i_* = 0 because *p_i_*_1_ = 1 and *a_i_* = 1. To get *π̅*, we then calculated the average value of *π_i_* across all *M* sites in the genome (including both variant and invariant sites):

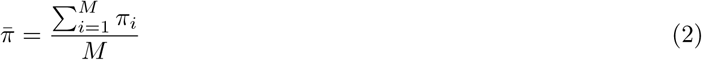

### *k*-mer counting

We chose to count *k*-mers of 30 base pairs (i.e. 30-mers) for all species in our dataset because previous *k*-mer-based analyses in plants typically analyzed *k*-mers in the range of 20 - 40 base pairs [Voichek and Weigel, 2020, Kim et al., 2020, VanWallendael and Alvarez, 2022, Ruperao et al., 2023] and because *k*-mers in this range can be reliably sequenced with short reads while capturing the majority of unique genomic sequences [Shajii et al., 2016, Ondov et al., 2016]. For each species, we built a database of the 30-mers that were present in the coding sequences of their reference genome using KMC (v3.2.1, Kokot et al. [2017]). Then, we counted 30-mers in each individuals’ sequencing reads using KMC, removing any 30-mers that matched the database of 30-mers found in its corresponding set of coding sequences. This step intended to focus our *k*-mers down to a set that is evolving more neutrally on average, analogously to how we focused on only 4-fold degenerate SNPs in our SNP-calling pipeline. The justification for this approach is that non-coding sequences generally have weaker signals of interspecies conservation compared to coding sequences [Woolfe et al., 2005, Siepel et al., 2005, Johnsson et al., 2014]. Although, similarly to 4-fold degenerate sites, many studies have observed non-coding sequences that appear to be under selective constraints [Margulies et al., 2003, Guo et al., 2007]. Thus, similar to the common analysis of 4-fold degenerate sites, our analysis is limited by an inability to completely remove the effects of selection on sequence diversity.

Although comparing our *k*-mer and nucleotide diversity metrics will be affected by differences between coding and non-coding sequences, many previous studies have found that the average diversity of non-coding regions is often very similar to average diversity at 4-fold degenerate sites [Moriyama and Powell, 1996, Maka-lowski and Boguski, 1998, Halushka et al., 1999, Zwick et al., 2000, Tenaillon et al., 2001, Nordborg et al., 2005, Branca et al., 2011, Williamson et al., 2014, Wang et al., 2016, Phung et al., 2016, Mattila et al., 2017]. Previous investigations of Lewontin’s paradox also found that diversity levels across species vary much more than diversity levels across different categories of putatively neutral sequences [Leffler et al., 2012, Buffalo, 2021] and subsequently pooled estimates of neutral diversity across different categories of sites. Similar to these previous studies, we thus assume that differences in linked selection between coding and non-coding sequences are negligible.

For most species in this study, we identified hundreds of millions of unique 30-mers. It would be computationally expensive to analyze all the 30-mers for every species. However, previous studies have shown that one can randomly downsample *k*-mer sets with very minimal effects on measures of genomic dissimilarity [Fofanov et al., 2004, Benoit et al., 2020]. Thus, we randomly downsampled each species’ 30-mer list to 10 million 30-mers with a frequency *≥* 5 in at least one sample in the species’ 30-mer list. The reason to include this frequency cut-off is to omit low frequency *k*-mers that result from sequencing errors [Ranallo-Benavidez et al., 2020]. We also chose to subset our 30-mer matrix to 10 million 30-mers to decrease disk space burden and because several previous studies show that subsets of only 1 million *k*-mers or less reliably estimate genetic dissimilarity in many systems [Ondov et al., 2016, Benoit et al., 2020, VanWallendael and Alvarez, 2022]. We then joined the subset *k*-mer counts for each individual into a single matrix for each species. We used this *k*-mer frequency matrix to measure genetic distance in two ways. First, we calculated Jaccard dissimilarity (*J_D_*, Ondov et al. [2016]) between each pair of individuals in a species’ dataset as:

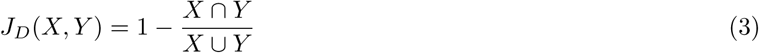

where *X* and *Y* represent sets of unique *k*-mers identified as present in two different read sets. A *k*-mer is defined as present if it’s frequency in a sample is *≥* 5, but cutoffs anywhere from 2 - 10 are commonly used in the literature [Voichek and Weigel, 2020, VanWallendael and Alvarez, 2022]. We used a frequency cutoff of 5 to make our workflow amenable to lower mean coverage datasets. To get the genome-wide average Jaccard dissimilarity (*J̅_D_*), we took the average of all the pairwise Jaccard dissimilarities.

Jaccard dissimilarity is likely the most commonly used *k*-mer-based diversity measure [Ondov et al., 2016]. However, whether a *k*-mer reaches the frequency threshold needed to be identified as preset in a sample depends on the sequencing depth for the sample [VanWallendael and Alvarez, 2022]. Thus, we also calculated Bray-Curtis dissimilarity (*B_D_*) between each pair of individuals in a species’ dataset as:

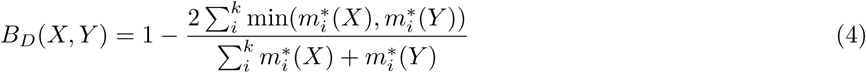

where 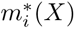 gives the normalized frequency of *k*-mer *i* in genome *X*. The normalized frequencies are calculated by taking each frequency *m_i_*(*X*) and dividing it by the sum of the raw frequencies as in Dubinkina et al. [2016]:

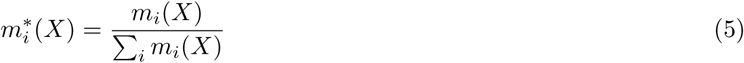

This step accounts for variation in coverage between samples on *k*-mer frequency. To get the genome-wide average Bray-Curtis dissimilarity (*B̅_D_*), we again took the average of all the pairwise Bray-Curtis dissimilarities. Note that both Jaccard and Bray-Curtis dissimilarity are scaled in their denominators by either the total number of unique *k*-mers or total number of *k*-mers respectively, analogous to how nucleotide diversity is scaled by the number of sites included in the calculation.

### Population size estimation

Following similar methods to Corbett-Detig et al. [2015] and Buffalo [2021], we defined current census population size (*N*) as the product of species range size (*R*) in square kilometers and population density (*D*) in individuals per square kilometer:

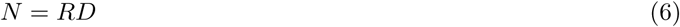

Estimation of both *R* and *D* are handled separately below. Importantly, these methods have the same drawback as described in Corbett-Detig et al. [2015] and Buffalo [2021]: contemporary estimates of *R* and *D* do not necessarily reflect the historical values of *R* and *D*. However, since nearly all the species in this study lack long-term historical data on their population size, it is not currently possible to estimate long-term historical *N* without making strong assumptions.

#### Range size estimation from GBIF occurrence data

We first estimated range size based on Global Biodiversity Information Facility (GBIF) occurrence data from the rgbif package [Chamberlain and Boettiger, 2017]. For each species, we identified its GBIF taxon key(s). If the species is domesticated, we used the taxon key(s) for a wild relative with an overlapping range when possible. We then downloaded all records associated with each taxon key that had an occurrence status of “PRESENT”, had coordinates that mapped to land, had any basis of record other than “FOSSIL SPECIMEN”, and recorded anywhere in a year *≥* 1943 and *≤* 2023. In addition, the records could not have any GBIF issue codes, except for the issue codes listed in Supplemental Methods. Similar to previous studies [Corbett-Detig et al., 2015, Buffalo, 2021], we estimated range size for domesticated species using GBIF occurrences from closely-related wild relatives because it is difficult to distinguish the native and introduced ranges of globally cultivated crop species with only occurrence data. Note, however, that we also used an additional method for estimating range size that is not burdened by this same assumption (see **Range size estimation from WCVP distribution maps**). The relatives used for each domesticated species is detailed in Table S1.

We followed methods of Buffalo [2021] to estimate range size from each species’ set of GBIF occurrence data using the package alphahull [Pateiro-Lopez and Rodriguez-Casal, 2022]. We started with splitting the occurrence data by continent, in order to avoid estimating ranges that overlapped with oceans. We also only kept occurrences with unique latitude-longitude values to reduce the computational burden of alphahull’s algorithms. We then added a small amount of random jitter (normally distributed with *µ* = 0 and *σ* = 1 *×* 10*^−^*^3^) to the latitude-longitude coordinates of each unique occurrence to avoid errors in the triangulation algorithm of alphahull, which can break when there are lots of colinear points. Finally, we filtered out any continents which had fewer than 20 unique occurrences of a species. The only exceptions to this rule were *Solanum stenotomum*, *Dioscorea alata*, and *Rhododendron griersonianum*, for which we only required 8, 6, and 3 occurrences respectively due to the rarity of these species and thus a paucity of occurrence data. We then used alphahull to compute the alpha shape of each continent subset, which can be thought of as the smallest possible convex shape that encloses a set of points in a plane. We defined the alpha parameter for the alphahull package to be 200. We then used the R packages sf [Pebesma, 2018] and rworldmap [South, 2011] to measure the sizes of the alpha shapes in square kilometers after projecting them onto the Earth’s surface. Finally, we took the estimated range polygons and filtered out ones that resided on continents in the introduced range of the species, as defined by the World Checklist of Vascular Plants (WCVP) [Govaerts et al., 2021]. The sum of the areas of the remaining polygons was our estimate of range size.

#### Range size estimation from WCVP distribution maps

We also estimated range size from expert-drawn species distribution maps instead of species occurrence data. We used the rWCVP package [Brown et al., 2023] to download distribution maps from WCVP [Govaerts et al., 2021]. We then estimated range size for each species as either (1) the sum of the areas of all map elements labeled as “native” or “extinct” for that species or (2) the sum of the areas of all map elements labeled as “native”, “invaded”, or “extinct” for that species. Regions with an occurrence label of “dubious” were excluded from downstream analyses. In contrast to GBIF-derived ranges, we used distribution maps for domesticated species in this estimate of range size because the maps discriminate between the native and introduced ranges of species.

#### Population density estimation from plant height

Similarly to previous studies, we use plant height as a proxy for plant population density [Corbett-Detig et al., 2015]. While it would be ideal to use actual population densities in our analyses, we could not find published estimates of population densities for many of the species in our dataset and all previous studies investigating Lewontin’s paradox rely on population size proxies [Leffler et al., 2012, Corbett-Detig et al., 2015, Filatov, 2019, Buffalo, 2021]. We elaborate further on the limitations of using proxies in the Discussion, but at the time of writing this manuscript using proxies is the only way to achieve a sufficient sample size for investigating Lewontin’s paradox.

We decided to use plant height rather than plant mass [Deng et al., 2012] as our measure of body size because plant height measurements are available for many more species in our dataset and also to make our results more comparable to previous studies that also use plant height [Corbett-Detig et al., 2015]. According to theory outlined in Deng et al. [2012], where *D* is population density, *M* is plant mass, and *h* is plant height, *D ∝ M^−^*^3*/*4^ and *M ∝ h*^8*/*3^. Combining these two relationships gives *D ∝* (*h*^8*/*3^)*^−^*^3*/*4^ which simplifies to *D ∝ h^−^*^2^. Adding this density-height relation to equation 6 gives our main proxy for population size:

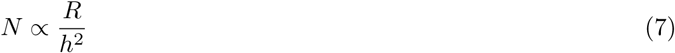

In our subsequent analyses, we refer to Equation 7 as the range size-squared height ratio and we convert *R* to square meters and *h* to meters to make the ratio unitless. As Equation 7 suggests, we do not expect the range size-squared height ratio to exactly equal the true population size or be interpretable as a number of individuals. Rather, it is a quantity we expect to scale with population size. To calculate the range size-squared height ratio for each species, we downloaded plant height data from the EOL, which mainly comprised records summarized from the TRY database. If no height measurements were available for a species in the EOL, then we used estimates we found in published scientific literature. The only exceptions to this were *Vanilla planifolia* and *Rhododendron griersonianum*, where our height estimates came from the Kew Botanical Gardens’ and the American Rhododendron Society’s websites, respectively. The sources used for each height value are cited in Table S1.

### Labeling species with genome size, mating system, ploidy, cultivation status, and life cycle habit

Table S1 contains citations for all studies that were used to label each species in our study with a genome size, mating system, ploidy level, cultivation status, and life-cycle habit. For determining genome size, we used estimates from flow cytometry and *k*-mer-spectra analyses whenever possible instead of using assembly size, since most assemblies do not contain the entire genome of the sequenced species. Most of our genome size estimates were 1C values acquired from publications cited in the Plant DNA C-values Database [Pellicer and Leitch, 2020]. Any estimates in terms of picograms (pg) of DNA were converted to base pairs using the following conversion factor: DNA in Mb = DNA in pg *×*0.978 *×* 10^9^ [Doležel et al., 2003]. If genome sizes in terms of pg were not available for a species, then we used the size of the species’ genome assembly as the genome size.

We next labeled each species with a mating system (selfing, outcrossing, mixed, or clonal), cultivation status (wild or cultivated), and life cycle habit (annual, biennial, perennial, or mixed) because previous studies showed these factors to be important determinants of diversity in plants [Chen et al., 2017]. For classifying species into different mating systems, we used methods similar to a previous study [Opedal et al., 2023] and generally considered species with outcrossing rate *<* 10 % as “selfing”, species with outcrossing rate between 10 - 90 % as “mixed”, and species with outcrossing rate *>* 90 % as “outcrossing” when estimates of outcrossing rates were available. In the absence of outcrossing rate data, we also labeled species described as generally self-incompatible as “outcrossing” and we labeled species described as selfing as “selfing”. The only exception to this was *Oryza brachyantha* for which we could not find mating system descriptions in peer-reviewed literature. Thus, we assumed that this species was most likely outcrossing because most of the other wild *Oryza* species in the dataset were classified as outcrossing. Because of the low number of mixed (14) and clonal (2) species in our dataset, we collapsed the selfing, mixed, and clonal species into a single “not outcrossing” category for later downstream analysis. Similarly, for life cycle habit, our dataset contained only 1 biennial species and 2 species that had a mixture of annual, biennial, and perennial forms. We combined these species with the perennial category to create a single “not annual” category. For cultivation status, we looked up each species in the EOL and classified species that had documented human uses (such as for food, fiber, fodder) or had some countries known to cultivate the species as “cultivated”. All other species that did not meet these criteria were classified as “wild”. The only exception to this was *Lactuca sativa*, which did not have any human uses listed in EOL at the time of writing this paper; however, it is commonly known as lettuce so we classified it as “cultivated”. Finally, for ploidy levels, when more than one cytotype was described as present within a species we labeled the species with it’s most common naturally-occurring cytotype. Citations to relevant literature used for each classification decision can be found in Table S1.

### Statistical analysis

The ultimate goal of our statistical analyses was to estimate the effect of our population size proxies on measures of diversity, comparing the effects of using *k*-mer-based or nucleotide diversity. To do this, we took an approach similar to Whitney et al. [2010] where we performed partial phylogenetic regressions controlling for evolutionary history (using a phylogeny obtained from timetree.org, Kumar et al. [2017, 2022]), mating system (outcrossing vs not outcrossing), cultivation status (wild vs cultivated), and life cycle habit (annual vs not annual). Similar to Whitney et al. [2010], we also scaled the dependent variables to be unitless with a mean of zero and unit variance across species (using the scale() function in R) before performing regression to make slopes more comparable across models and account for the inherent differences in unit between nucleotide and *k*-mer diversity metrics. This approach can be summarized as follows:

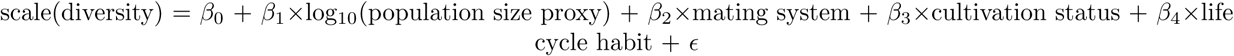

where population size proxy refers to either Equation 7 or it’s components (range size and plant height), covariance in the residuals is given by *V ar*[*ɛ*] = Ω, diversity was estimated using either SNPs (log_10_(*π̅*)) or *k*-mers (*J̅_D_* or *B̅_D_*), and the scale() function performs a z-transformation to make diversity unitless with mean of zero and unit variance. We also constructed a separate set of models where we included genome size as a covariate:

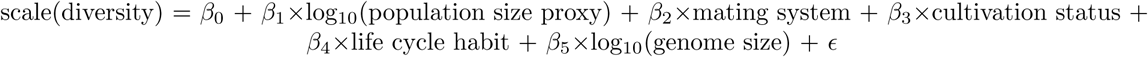

We controlled for genome size in a separate set of models because we had conflicting expectations on whether genome size would be a confounder or a mediator of the population size-diversity relationship. In other words, the effect of population size on diversity could act through genome size, since small populations may not experience strong enough selection to purge deleterious insertions [Lynch and Conery, 2003]. Including genome size as a covariate in this case would artificially diminish the estimated effect of population size on diversity. Alternatively, genome size could fundamentally alter the mode of adaptation in plant species [Mei et al., 2018], making genome size a confounder of the population size-diversity relationship.

After constructing our models, we visualized the relationship between population size and diversity or genome size and diversity with partial regression plots, following methods from Riddell [1977] and Blomberg et al. [2012]. Beginning with our initial phylogenetic least squares model:

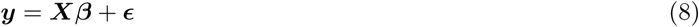

where ***y*** is a vector of diversity values, *X* is the design matrix, ***β*** is a vector of regression coefficients, and ***ɛ*** is a vector of residuals distributed normally about 0 with phylogenetic variance-covariance matrix Ω. Using the variance-covariance matrix output from the caper R package [Orme et al., 2018], we first performed Cholesky decomposition to get matrix ***C*** such that:

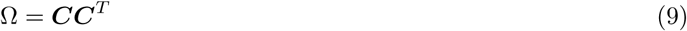

We then took the inverse matrix ***C****^−^*^1^ and left-multiplied both sides of our regression equations to get:

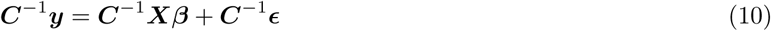

Which we will rewrite as:

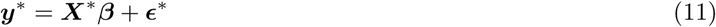

where ***y****^∗^* = ***C****^−^*^1^***y***, ***X****^∗^* = ***C****^−^*^1^***X***, and ***ɛ****^∗^* = ***C****^−^*^1^***ɛ***. In vector form, this equation is now:

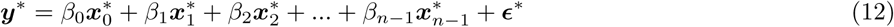

where 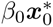 is our intercept (Note that ***x*_0_** was initially a column of 1’s before being transformed by ***C****^−^*^1^). After fitting this model to our data with the standard lm() function in R, we collected all terms besides the primary variable of interest, 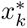 (which would be a population size proxy or genome size in our case), and subtracted them from both sides of the equation to get:

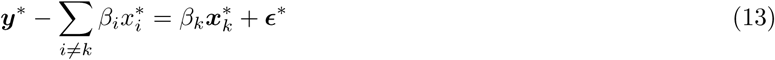

We then plotted the values of 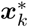 against 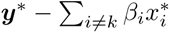, interpreting the slope as the effect of the primary variable on the response, scaled for phylogenetic relationships and adjusted for the effects of confounding factors.

## Results

### Low diversity species explained by low mean coverage

In total, we processed *>*205 terabases of publicly available sequencing data from the SRA over approximately 12 months of wall time, split between a maximum of 512 cores and 50 TB of disk space. There were 112 species in our initial dataset, each with estimates of population size proxies, nucleotide diversity, and *k*-mer diversity (Fig. 1). Out of these 112 species, 102 were diploids, 9 were tetraploids, and one was hexaploid, with haploid genome sizes ranging from 105 Mb to 5.06 Gb (Table S1). These species were further broken down into 57 annual species vs 55 not annual species (which were predominately perennial), 31 wild vs 81 cultivated species, and 55 outcrossing vs 57 not outcrossing species (which were predominantly selfing). Species classified as annual also tended to not be classified as outcrossing (*χ*^2^ = 18.9, p = 1.4 *×* 10*^−^*^5^, Fig. S1C). However, cultivation status was independent of both life cycle habit (*χ*^2^ = 4.07 *×* 10*^−^*^31^, p = 1, Fig. S1A) and mating system (*χ*^2^ = 0.53, p = 0.47, Fig. S1B). There were no missing values for any of the variables investigated in this study, but there were three species with zero variant sites called that we omitted from all downstream analyses.

**Figure 1.**
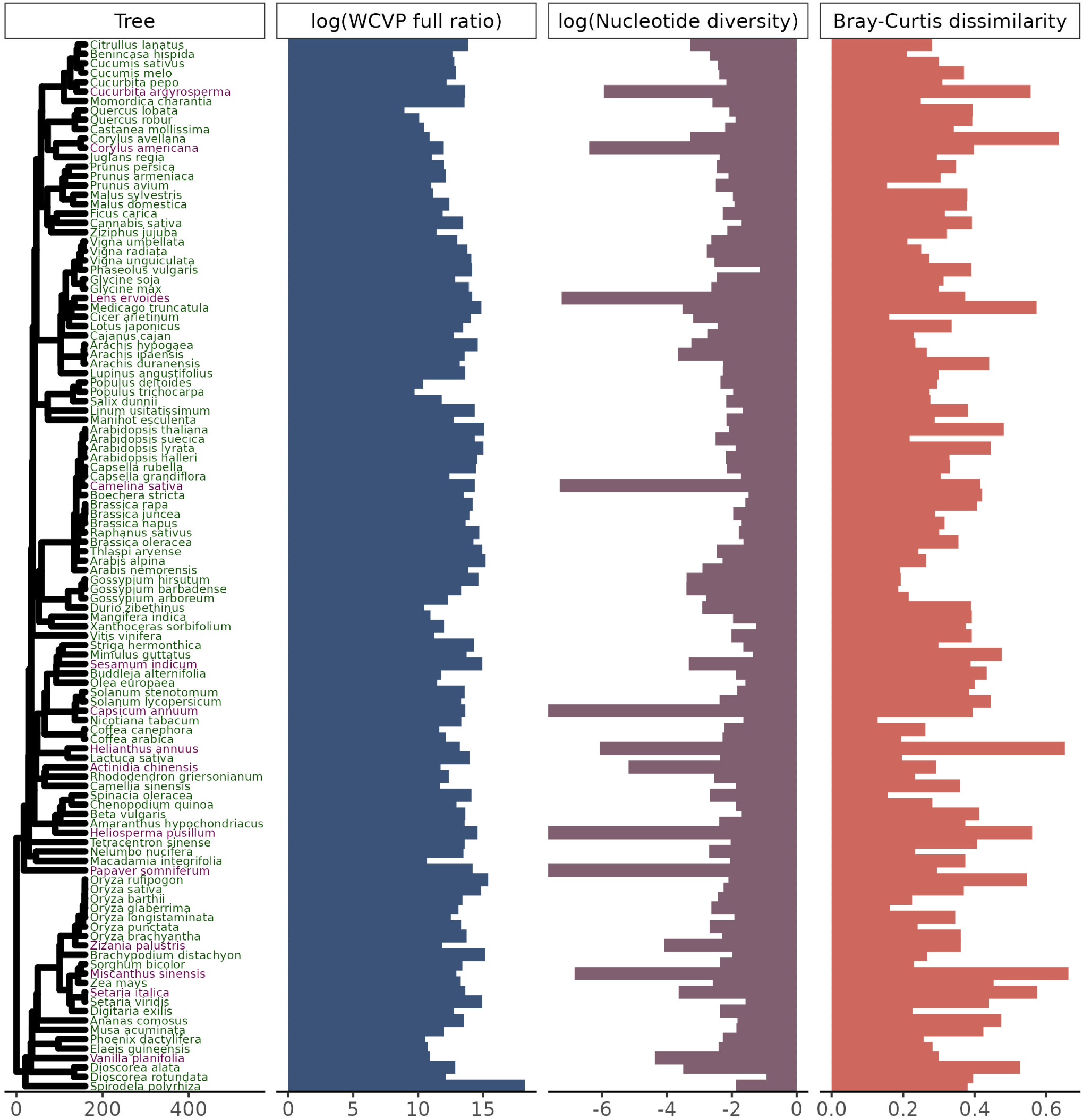
Our study includes 112 plant species across a wide range of population sizes and diversity levels. Species labeled in purple were considered outliers and omitted from downstream analyses (see Fig. 2), but species labeled in green were retained. The phylogenetic tree is scaled in millions of years. The WCVP full ratio is a unitless population size proxy equal to the ratio of range area, estimated using WCVP range maps, to squared plant height and is log-transformed (base 10). Nucleotide diversity is genome-wide average diversity at four-fold degenerate sites, log-transformed (base 10). *Capsicum annuum*, *Heliosperma pusillum*, and *Papaver somniferum* had nucleotide diversity values of zero and so have bars at the plotting limit (log(0) = *−∞*). Bray-Curtis dissimilarity is average pairwise Bray-Curtis dissimilarity across all pairs of individuals in a species’ sample.

Before testing our central hypothesis, we investigated whether technical sequencing variables could explain any of the diversity values observed in our dataset. As would be expected for a meta-analysis of previously published data, sequencing parameters varied between species. The number of individuals sampled in each species varied from 3 to 1200 and the average depth of sequencing per individual varied from 0.028x to 79.7x (Fig. S2). Variation in the depth of sequencing between individuals, quantified as the coefficient of variation in base pairs sequenced, varied about 50-fold from 0.030 to 1.6 (Fig. S2). Mean coverage correlated with both nucleotide diversity (*ρ* = 0.33, p = 0.00033, Fig. S3A) and *k*-mer diversity (Jaccard: *ρ*= −0.53, p = 2.6*×*10*^−^*^9^, Fig. S3D; Bray-Curtis: *ρ* = −0.34, p = 0.00021, Fig. S3G). Coefficient of variation in bp sequenced correlated strongly with *k*-mer diversity (Jaccard: *ρ* = 0.36, p = 0.00013, Fig. S3E; Bray-Curtis: *ρ* = 0.42, p = 4.7 *×* 10*^−^*^6^, Fig. S3H) but not nucleotide diversity (*ρ* = −0.088, p = 0.36, Fig. S3B). The number of individuals sequenced did not correlate with either nucleotide diversity or *k*-mer diversity (Fig. S3C, S3F, S3I).

While screening the data for outliers, we expected that nucleotide diversity and *k*-mer-based diversity would be positively correlated across species and that deviations from this expectation might result from technical variation in how sequencing was performed. Overall, we observed that species with lower coverage did not follow the expected positive relationship between nucleotide and *k*-mer diversity (Fig. 2A, Fig. S4A). In contrast, there was no clear pattern in how the coefficient of variation in base pairs sequenced (Fig. S5) or the number of individuals sequenced (Fig. S6) affected the correlation between *k*-mer dissimilarity and nucleotide diversity. Based on these results, we removed 10 species from our dataset with mean coverage per individual *≤* 0.5x as well as 4 species with higher coverage but fewer than 1000 variant sites called. This included three species (*Capsicum annuum*, *Heliosperma pusillum*, and *Papaver somniferum*) with zero variant sites called. The correlation between nucleotide diversity and *k*-mer diversity was much more significant after excluding these species (Jaccard: *ρ* = 0.34, p = 0.00068, Fig. S4B; Bray-Curtis: *ρ* = 0.49, p = 3.6*×*10*^−^*^7^, Fig. 2B). In total, we kept data for 98 species for downstream hypothesis testing.

**Figure 2.**
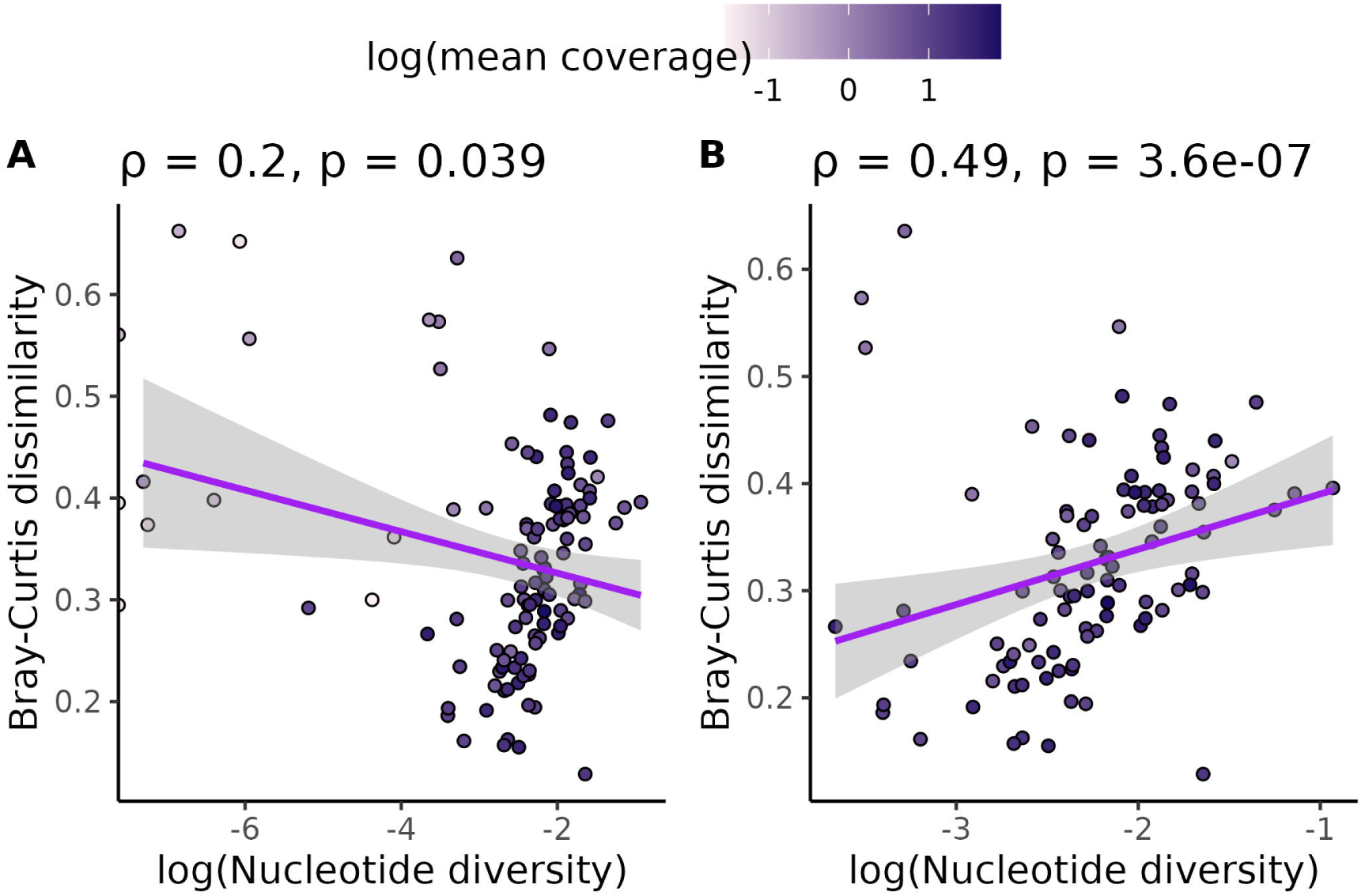
Omitting species with low coverage and low numbers of variant calls increased the positive correlation between nucleotide and *k*-mer diversity. (A) shows the relationship between *k*-mer diversity and nucleotide diversity without omitting species with *≤* 0.5x coverage or *≤* 1000 SNP calls. (B) shows the same relationship, except species with *≤* 0.5x coverage or *≤* 1000 SNP calls are omitted. Each data point is a species. All species’ points are colored by the log (base 10) of average genome-wide coverage per individual for that species. Purple lines are linear regressions with 95% confidence intervals shaded in gray. Values across the top of each plot are Spearman correlation coefficients (*ρ*) and p-values that test whether each correlation coefficient differs from zero.

### Range size-squared height ratio varies over more orders of magnitude than nucleotide diversity

We next investigated whether Lewontin’s paradox applied to our dataset by comparing diversity estimates against population size proxies. For each species, we estimated range size using either GBIF occurrence data or WCVP range maps. Estimates from these two methods were significantly correlated no matter whether invaded ranges (as defined in the WCVP range maps) were included (*ρ* = 0.31, p = 0.00096, Fig. S7A) or excluded (*ρ* = 0.48, p = 7.3*×*10*^−^*^8^, Fig S7B). The omission of invaded ranges lowered the range size of several plant species based on WCVP range maps (Fig. S7C) but had less effect on ranges estimated from GBIF occurrence data (Fig. S7D).

We then calculated the ratio of range size to squared plant height (Equation 7) using height values from the EOL. We used this ratio as our primary population size proxy in downstream analyses. After excluding species with *<* 0.5x coverage and *<* 1000 variant sites called (Fig. 2), nucleotide diversity varied over about 4 orders of magnitude for the species in our dataset (from 0.00021 to 0.117, Table S2), while the ratio of range size to squared plant height based on WCVP and GBIF range estimation methods (including both native and invaded ranges) varied over 10 (from 8.9 *×* 10^8^ to 1.7 *×* 10^18^) and 13 (from 8.6 *×* 10^5^ to 1.5 *×* 10^18^) orders of magnitude, respectively (Table S2). Mean pairwise Bray-Curtis dissimilarity values varied about 4.9-fold across species, from 0.13 to 0.64, while mean pairwise Jaccard dissimilarity varied about 22-fold, from 0.040 to 0.87 (Table S2). Bray-Curtis dissimilarity values correlated with Jaccard dissimilarity values across species (*ρ* = 0.76, p *<* 2.2 *×* 10*^−^*^16^, Fig. S8).

### *k*-mer diversity scales with population size proxies more than nucleotide diversity

The core of Lewontin’s paradox is that a population’s diversity does not scale much with population size. If *k*-mers capture a wider range of genetic variation compared to SNPs, population size will scale more with *k*-mer diversity than nucleotide diversity. If we did not control for shared evolutionary history or any confounding variables (mating system, life cycle habit, cultivation status, or genome size), then none of our diversity measures significantly correlated with the range size-squared height ratio (Fig. S9). After controlling for confounding variables, nucleotide diversity marginally scaled with the range size-squared height ratio (*β* = 0.14, SE = 0.056, p = 0.017, Fig. S10A). However, the relationship between *k*-mer diversity and the range size-squared height ratio was highly significant, with generally a greater slope (Jaccard: *β* = 0.64, SE = 0.096 p = 2.2 *×* 10*^−^*^9^, Fig. S10B; Bray-Curtis dissimilarity: *β* =0.79, SE = 0.11, p = 7.3 *×* 10*^−^*^11^, Fig. S10C). We observed the same qualitative trend when we included both native and invaded ranges in the range size-squared height ratio (Fig. S10D-F), or used the GBIF-based range estimates instead of WCVP-based estimates (Fig. S11). Interestingly, we often observed Bray-Curtis dissimilarity having a larger slope with the range size-squared height ratio compared to Jaccard dissimilarity (*β* = 0.64 vs 0.79 Fig. S10B-C), but models where Bray-Curtis dissimilarity was the response variable generally had lower adjusted *R*^2^ (Table S4).

We also analyzed range size and plant height separately as population size proxies (Fig. S12-S14). Overall, WCVP-estimated range size significantly affected nucleotide diversity (*β* = 0.29, SE = 0.072, p = 0.00011, Fig. S12A) and *k*-mer diversity (Jaccard: *β* = 0.92, SE = 0.13, p = 9.9*×*10*^−^*^11^, Fig. S12B; Bray-Curtis: *β* = 1.2, SE = 0.13,p = 3.2*×*10*^−^*^14^, Fig. S12C), and this trend held when we estimated range size from GBIF occurrences (Fig. S13A-C) or included invaded range area (Fig. S12D-F and Fig. S13D-F). On the other hand, plant height did not scale with nucleotide diversity (*β* = 0.13, SE = 0.19, p = 0.5, Fig. S14A), but marginally scaled downward with increasing *k*-mer diversity (Jaccard: *β* = *−*0.78, SE =0.38, p = 0.046, Fig. 14B; Bray-Curtis: *β* = *−*0.77, SE = 0.44, p = 0.088, Fig. S14C).

Finally, we repeated our partial phylogenetic regressions controlling for genome size as an additional covariate. In this case, nucleotide diversity did not scale with the range size-squared height ratio (*β* = 0.035, SE = 0.063, p = 0.58, Fig. 3A), but *k*-mer diversity did (Jaccard: *β* = 0.54, SE = 0.093, p = 8.8 *×* 10*^−^*^8^, Fig. S15; Bray-Curtis: *β* = 0.7, SE = 0.098, p = 2.2*×*10*^−^*^10^, Fig. 3B). Again, we got qualitatively similar results when we excluded invaded ranges in our range size estimates (Fig. S16), used GBIF occurrences to estimate range size-squared height ratio (Fig. S17) or used WCVP range size as the population size proxy (Fig. S18). However, GBIF range size by itself did not scale with Jaccard dissimilarity (Fig. S19B, S19E). Increased plant height associated with decreased *k*-mer diversity, but had no significant relationship with nucleotide diversity (Fig. S20).

**Figure 3.**
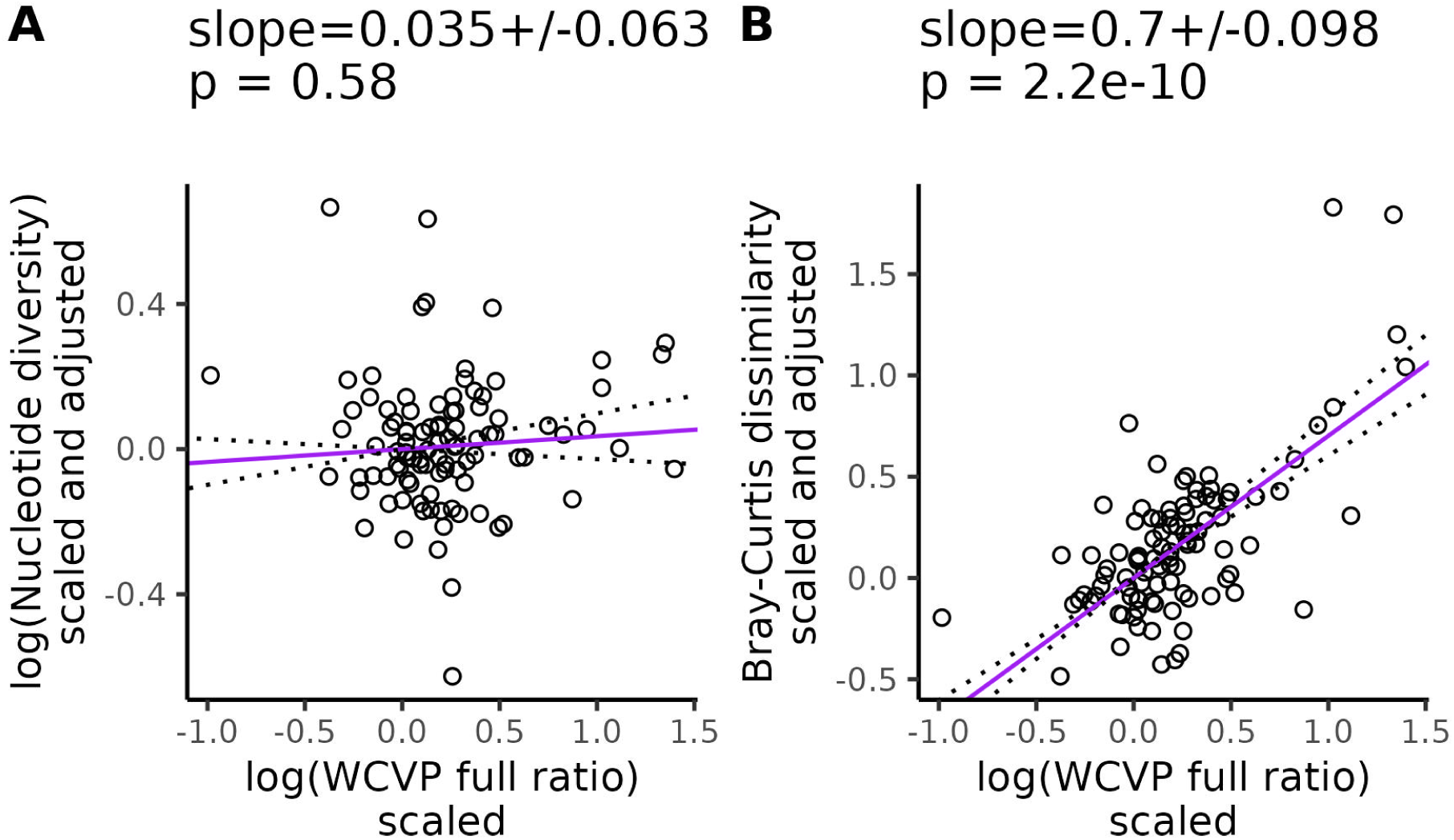
*k*-mer diversity scales with population size proxies after controlling for genome size, life cycle habit, mating system, and cultivation status. WCVP full ratio is a population size proxy estimated as the ratio of range size recorded in WCVP range maps (including invaded ranges) to squared plant height. Purple lines are partial phylogenetic regression lines between diversity levels and the population size proxy (see Equation 13) after scaling diversity levels to a standard normal distribution (mean = 0, variance = 1), followed by scaling diversity levels and population sizes according to their phylogenetic relatedness, and finally adjusting for the confounding variables (genome size, life cycle habit, mating system, and cultivation status). The values at the top of each plot give the slope of the partial regression *±* one standard error and p-values testing whether the slopes differ from zero. Dotted lines show the partial regression slope *±* one standard error.

### *k*-mer diversity scales with genome size more than nucleotide diversity

We also investigated the relationship between diversity and genome size because we expected genome size to potentially play a role in the mechanism underlying the greater scaling of *k*-mer diversity with population size. Genome size is often a strong predictor of diversity [Lynch and Conery, 2003]. Among eukaryotes, variation in genome size is largely explained by variation in transposable element abundance [Flavell et al., 1974, Kidwell, 2002, Lynch and Conery, 2003, Muñoz-Diez et al., 2012, Tenaillon et al., 2011, Nystedt et al., 2013, Ibarra-Laclette et al., 2013], which contribute substantially to the repetitive sequence content of genomes and increase the difficulty of aligning short reads to a reference genome (reviewed in Goerner-Potvin and Bourque [2018]). Thus, our expectation was that *k*-mer-based diversity measures are more sensitive to genome size variation compared to nucleotide diversity.

Increasing genome size was associated with decreasing *k*-mer diversity (Jaccard: *β* = *−*3.7, SE = 0.42, p = 8.4*×*10*^−^*^14^, Fig. S21; Bray-Curtis: *β* = *−*4.2, SE = 0.45, p = 4.5*×*10*^−^*^15^, Fig. 4B) and nucleotide diversity (*β −* 1.8, SE = 0.29, p = 1.4 *×* 10*^−^*^8^, Fig. 4A), after controlling for variation in the range size-squared height ratio, mating system, life cycle habit, cultivation status, and evolutionary history. We got qualitatively similar results when the population size proxy we corrected for excluded invaded ranges (Fig. S22), or if our population size proxy was based on GBIF occurrences (Fig. S23), or we used range size or plant height individually to control for population size variation (Fig. S24-S26). Across all of these analyses, the partial regression relationship between genome size and diversity was always significantly negative.

**Figure 4.**
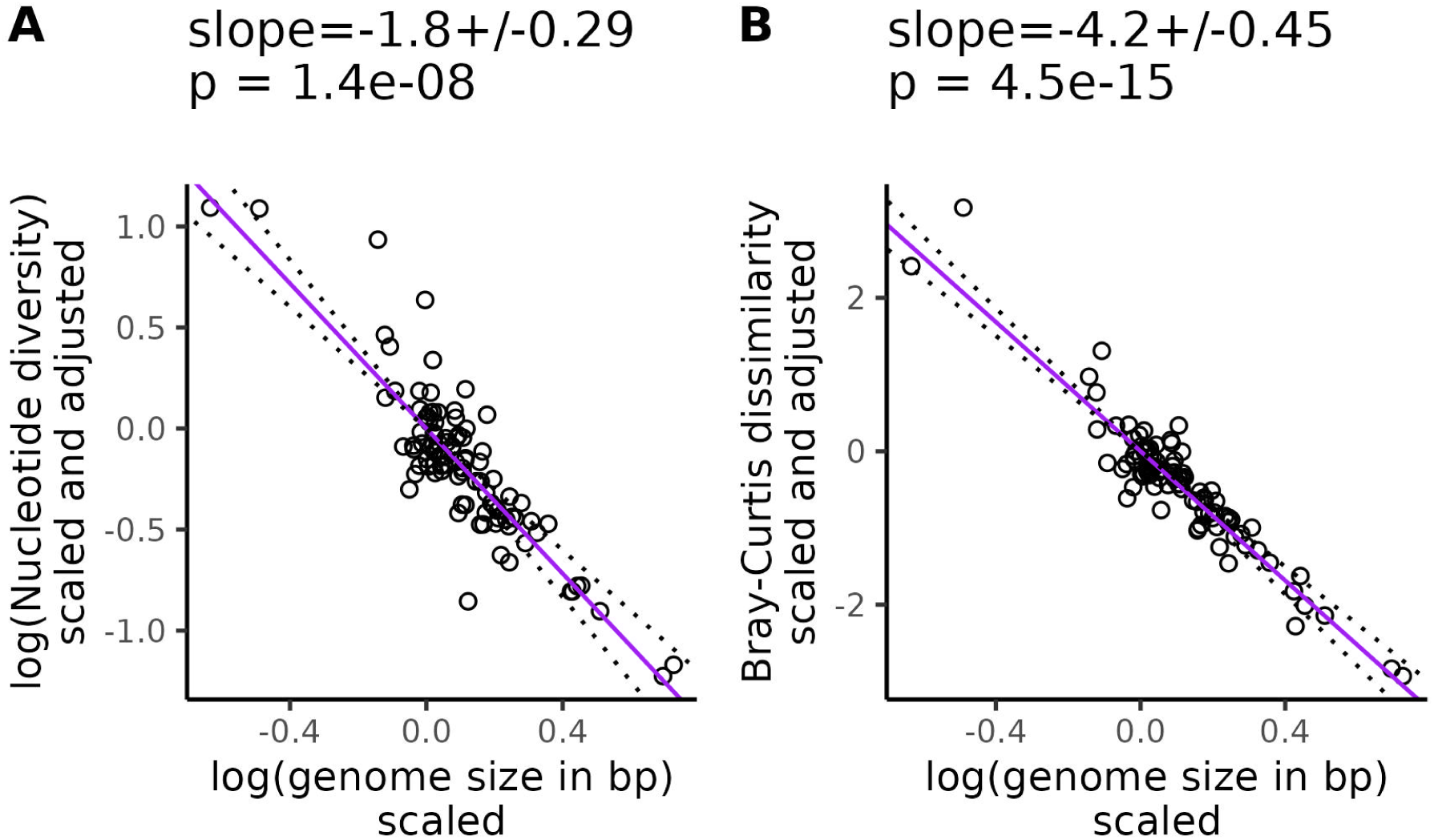
*k*-mer diversity is more sensitive to genome size variation than nucleotide diversity. Purple lines are partial phylogenetic regression lines between diversity levels and genome size (see Equation 13) after scaling diversity levels to a standard normal distribution (mean = 0, variance = 1), followed by scaling diversity levels and population sizes according to their phylogenetic relatedness, and finally adjusting for the confounding effects of mating system, cultivation status, life cycle habit, and population size. Here we used the ratio of range size to squared plant height, where range size was estimated from ranges in WCVP range maps (including invaded ranges). The values at the top of each plot give the slope of the partial regression *±* one standard error and p-values testing whether the slopes differ from zero. Dotted lines show the partial regression slope *±* one standard error.

## Discussion

Our primary goal was to investigate whether genomic approaches that can capture more genetic variation than standard SNP-based methods can explain the longstanding observation that species with large population sizes have less genetic variation than expected. After careful accounting for potential technical and phylogenetic confounding, the slope between *k*-mer-based diversity and the range size-squared height ratio was up to 20 times larger than the same slope for nucleotide diversity (*β* = 0.035 vs 0.7, Fig. 3). We observed similar results across the two different measures of range size (Fig. S17) and *k*-mer diversity (Fig. S15). We also observed that *k*-mer-based diversity is more sensitive to variation in genome size compared to nucleotide diversity (Fig. 4). Overall, these results suggests that diversity missed by SNPs explains part of Lewontin’s paradox in plants, consistent with literature suggesting that SNPs provide an incomplete picture of genome-wide polymorphism [Schmidt et al., 2021, VanWallendael and Alvarez, 2022, Jaegle et al., 2023, Sopniewski and Catullo, 2024].

One limitation of our investigation was that we were not able to compare our *k*-mer diversity scales to a neutral expectation of how *k*-mer diversity scales with *N_e_*. Doing so would have allowed us to estimate what proportion of Lewontin’s paradox is explained by using *k*-mer diversity instead of nucleotide diversity measures. Instead we can only compare the slopes of how *k*-mer and nucleotide diversity scale with population size proxies. We deliberately avoided comparing our data to a neutral expectation for two main reasons. First, we can only estimate proxies of census size that are not interpretable as numbers of individuals, which is what a theoretical expectation would most likely be based on. Furthermore, robustly estimating the diversity-census size relationship across species requires controlling for evolutionary history and other confounding variables. This will transform the axes of a diversity-population size partial regression plot into a scale that’s not interpretable in the units of the original measures (see Equation 10). Thus, we must restrict our conclusions to whether *k*-mer diversity scales with census size proxies faster than nucleotide diversity. This observation is consistent with the hypothesis that the exclusion of non-reference variation explains a part of Lewontin’s paradox. However, exactly what proportion of the paradox is explained by our results remains unknown.

As with all regression-based analyses, our results are also ultimately sensitive to error in the measurement of both covariates (population size proxies, genome size, mating system, life cycle habit, or cultivation status) and outcome variables (nucleotide or *k*-mer diversity). Random covariate measurement errors (i.e. error that is not systematically higher/lower for different values of the covariate) bias regression coefficients toward zero [Hutcheon et al., 2010, Nab et al., 2021]. Similarly, random measurement error in the outcome variables increases the standard errors of the covariates, weakening the statistical significance of detected relationships [Hutcheon et al., 2010]. However, our results remain statistically significant despite the potential for error. Our study is also unique in the multiple steps we took to limit the influence of systematic measurement errors on our coefficients. First, we reanalyzed all population-level sequencing data with a single pipeline to limit between-study variation and the impact of bioinformatic parameter choices on our analysis [Mirchandani et al., 2024]. Second, to minimize error in both nucleotide and *k*-mer diversity measures, we omitted species with coverage below 0.5x from our study, because having low coverage strongly correlated with having low diversity (Fig. 2). This threshold is consistent with previous *k*-mer-based phylogenetic studies that found dropping coverage to 0.5x changes tree topologies compared to coverage levels *≥* 1x [Sandell et al., 2022]. Third, we accounted for the presence of missing data in calculations of nucleotide diversity [Schmidt et al., 2021, Korunes and Samuk, 2021]. And finally, we estimated range size with two different methods (WCVP range maps and GBIF occurrence records, Fig. S6). Although we could not control for some covariates [Willis, 1922, Romiguier et al., 2014, Guo et al., 2024] due to a dearth of data, our study is still the largest reanalysis of population-level sequencing data in plants that we know of to date. The availability of our workflow also makes it easy for our study to be extended as more population-level sequencing data is released.

Another limitation of most investigations into Lewontin’s paradox is the assumption that contemporary population size estimates are good proxies for historic population sizes [Corbett-Detig et al., 2015, Buffalo and Coop, 2020]. While the long-term harmonic mean of the effective population size determines diversity levels within a population [Wright, 1940], population size proxies such as range size and plant height only reflect the current census population size of a species. The separation of plant range maps into native and invaded ranges [Brown et al., 2023] offered an opportunity to test the robustness of our results to invasion-related range size changes. Overall, our observations were remarkably similar no matter whether we included or excluded invaded ranges in our population size proxies (Fig. S17A-C vs Fig. S17D-F). Part of this apparent robustness was due to the insensitivity of our GBIF-based range size estimates to the inclusion of invaded ranges (Fig. S7D). However, our WCVP-based range size estimates were drastically altered by the inclusion of invaded ranges (Fig. S7C) and still yielded similar results (Fig. 3, S15, S16). Although we cannot rule out the possibility that older historical events have affected contemporary diversity levels, our results appear to be robust to some recent human-caused population size changes.

Interestingly, the estimated effect of our population size proxies on diversity was often slightly larger for Bray-Curtis dissimilarity than Jaccard dissimilarity (for example, *β* = 0.7 vs 0.54 from Fig. 3B vs Fig. S15, Table S4). In contrast, the range size-squared height ratio was often slightly more predictive of Jaccard dissimilarity than Bray-Curtis dissimilarity (Table S4). We could not test whether these trends were statistically significant, but the benefits of different *k*-mer metrics in predicting measures of population size warrant further study. Our expectation is that *k*-mer diversity measures based on frequency, such as Bray-Curtis dissimilarity, better capture diversity compared to measures based on purely *k*-mer presence/absence, such as Jaccard dissimilarity, because they explicitly measure copy number variation. However, accurately measuring *k*-mer frequencies likely requires higher sequencing coverage than calling presence/absence, which could explain why Bray-Curtis dissimilarity generally scaled more with population size but had a lower *R*^2^ compared to Jaccard dissimilarity (Table S4). Future studies using higher coverage population level sequencing data could help test this hypothesis.

*k*-mer frequencies are known to be highly informative of genomic structure, with one common application of *k*-mers being the estimation of genome size [Vurture et al., 2017, Pflug et al., 2020]. Similar to previous studies, we observed that nucleotide diversity was negatively correlated with genome size [Lynch and Conery, 2003, Chen et al., 2017], but we observed an even stronger negative correlation for *k*-mer diversity (*β* = *−*1.8, SE = 0.29 vs *β* = −3.7, SE = 0.42 in Fig. 4). *k*-mers also appeared to explain diversity patterns that scaled with population size beyond those explained by genome size, while nucleotide diversity did not. After controlling for genome size, the relationship between our population size proxies and nucleotide diversity was not significant (Fig. 3A, S17-S19 panels A and D), but the relationship between *k*-mer diversity and population size proxies was often still highly significant (Fig. 3B, S17-S19 panels B; C; E; F). The only exception was that Jaccard dissimilarity did not significantly scale with GBIF-based estimates of range size (Fig. S19B, S19E). This additional scaling of *k*-mer diversity with population size beyond just the effects of genome size and confounding variables suggests that *k*-mers capture some element of the population size-diversity relationship that is absent from nucleotide diversity.

Our results do not negate the fact that other important factors also underlie Lewontin’s paradox, such as past demographic fluctuations and linked selection. However, our results do suggest that future studies of Lewontin’s paradox would benefit from considering diversity outside one reference genome. The increasing availability of pangenomes across species [Göktay et al., 2021, Zhou et al., 2022, Rice et al., 2023, Wang et al., 2023] offers many opportunities to revisit this classic population genetics question. While our results suggest that including non-reference variation may partially satisfy Lewontin’s paradox, exactly how much of the paradox is explained by non-reference variation, whether our findings apply outside of plants, and the relative importance of non-reference variation to other factors in explaining Lewontin’s paradox is still unknown. Ideal future studies would use pangenomic genotyping methods across a wide range of species with a standardized pipeline, combined with multiple proxies of population size. Altogether, these methodological developments will hopefully reveal a more wholistic picture of variation across the tree of life.

## Data availability

Our entire analysis is packaged as a snakemake workflow stored here: https://github.com/milesroberts-123/tajimasDacrossSpecies. Table S1 contains the metadata for all of the datasets used in this study, including sources for genome assemblies, genome annotations, population-level sequencing datasets, and GBIF observations. Table S2 contains all of the covariate and response variable values used for fitting our phylogenetic least squares models. Table S3 contains the estimated coefficients of all of our phylogenetic least squares models and their related statistics, including p-values and standard errors. Table S4 contains the model-level statistics for each phylogenetic least squares model, including *R*^2^ values and F-test results. If necessary, we are also prepared to publish the following datasets upon acceptance of this manuscript in the accepting journal’s preferred repository: matrices of *k*-mer counts (93 G), VCF files of filtered variants (202 G), multiqc reports of fastp read trimming (244 M), species range maps (87M, downloaded from Plants of the World Online), and plant height values (downloaded from Encyclopedia of Life), and our species tree (downloaded from timetree.org).

## Acknowledgments

We would like to thank Jeff Conner, Husain Agha, Sophie Buysse, Nathan Catlin, Adrian Platts, Gabrielle Sandstedt and the rest of the Josephs lab for informal comments on early drafts of this manuscript. We would also like to thank the Institute for Cyber-Enabled Research at Michigan State University for providing the computing power used for this research. Finally, we would like to thank the hundreds of scientists throughout the world that sequenced plant populations and publicly released their raw data. Without them this work would not have been possible.

## Author contributions

Both M.D.R and E.B.J contributed to the initial conceptualization and planning of this research. M.D.R wrote all of the code and conducted all statistical analyses. Both M.D.R and E.B.J contributed to drafting, editing, and reviewing the manuscript.

