## Supplemental methods and figures for "Previously unmeasured genetic diversity explains part of Lewontin’s paradox in a *k*-mer-based meta-analysis of 112 plant species"

2024-09-08

#### Contents

**1 Supplemental methods** **2**

**2 Supplemental figures: Exploring relationships in data before outlier removal** **3**

**3 Supplemental figures: Population size proxy vs diversity relationships after controlling for phylogeny and life-history variables, but not controlling for genome size** **12**

**4 Supplemental figures: Population size proxy vs diversity relationships after controlling for phylogeny, life-history variables, and genome size** **17**

**5 Supplemental figures: Genome size vs diversity relationships after controlling for population size proxies, phylogeny, and life-history variables** **23**

### 1 Supplemental methods

The records that we used to estimate range size from GBIF occurrence data could not have any issue codes except for the following:

- "AMBIGUOUS_COLLECTION"
- "AMBIGUOUS_INSTITUTION"
- "COLLECTION_MATCH_FUZZY"
- "COLLECTION_MATCH_NONE"
- "CONTINENT_DERIVED_FROM_COORDINATES"
- "COORDINATE_ROUNDED"
- "COUNTRY_DERIVED_FROM_COORDINATES"
- "COUNTRY_MISMATCH"
- "DEPTH_MIN_MAX_SWAPPED"
- "DEPTH_NON_NUMERIC"
- "DEPTH_NOT_METRIC"
- "DEPTH_UNLIKELY"
- "DIFFERENT_OWNER_INSTITUTION"
- "ELEVATION_MIN_MAX_SWAPPED"
- "ELEVATION_NON_NUMERIC"
- "ELEVATION_NOT_METRIC"
- "ELEVATION_UNLIKELY"
- "GEODETIC_DATUM_ASSUMED_WGS84"
- "INSTITUTION_COLLECTION_MISMATCH"
- "INSTITUTION_MATCH_FUZZY"
- "INSTITUTION_MATCH_NONE"
- "OCCURRENCE_STATUS_INFERRED_FROM_BASIS_OF_RECORD"
- "OCCURRENCE_STATUS_INFERRED_FROM_INDIVIDUAL_COUNT"

#### 2 Supplemental figures: Exploring relationships in data before outlier removal

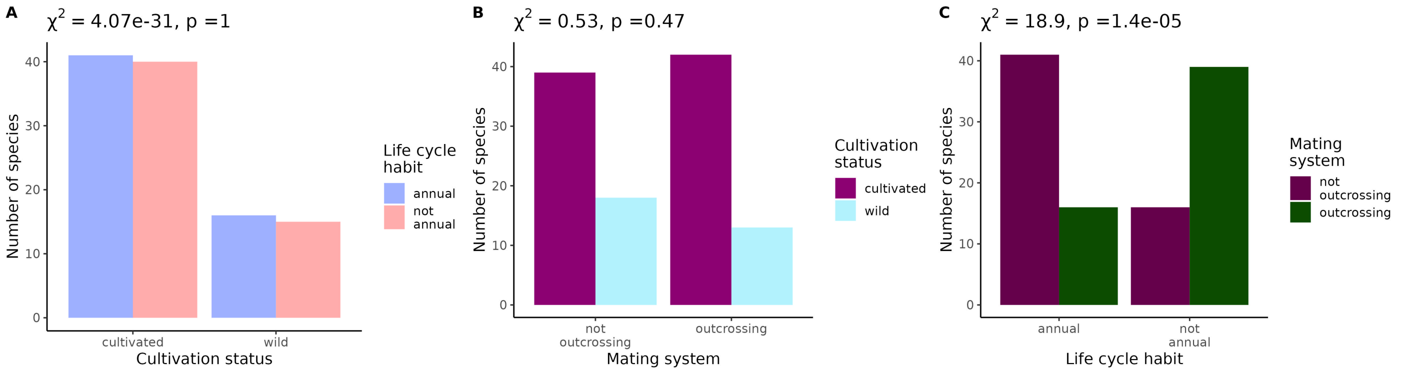


Figure S1: **Tests of independence between life-history traits included in this study.** Values across the top of each plot give the results of a $\chi^2$ test of independence between each pair of life-history traits.

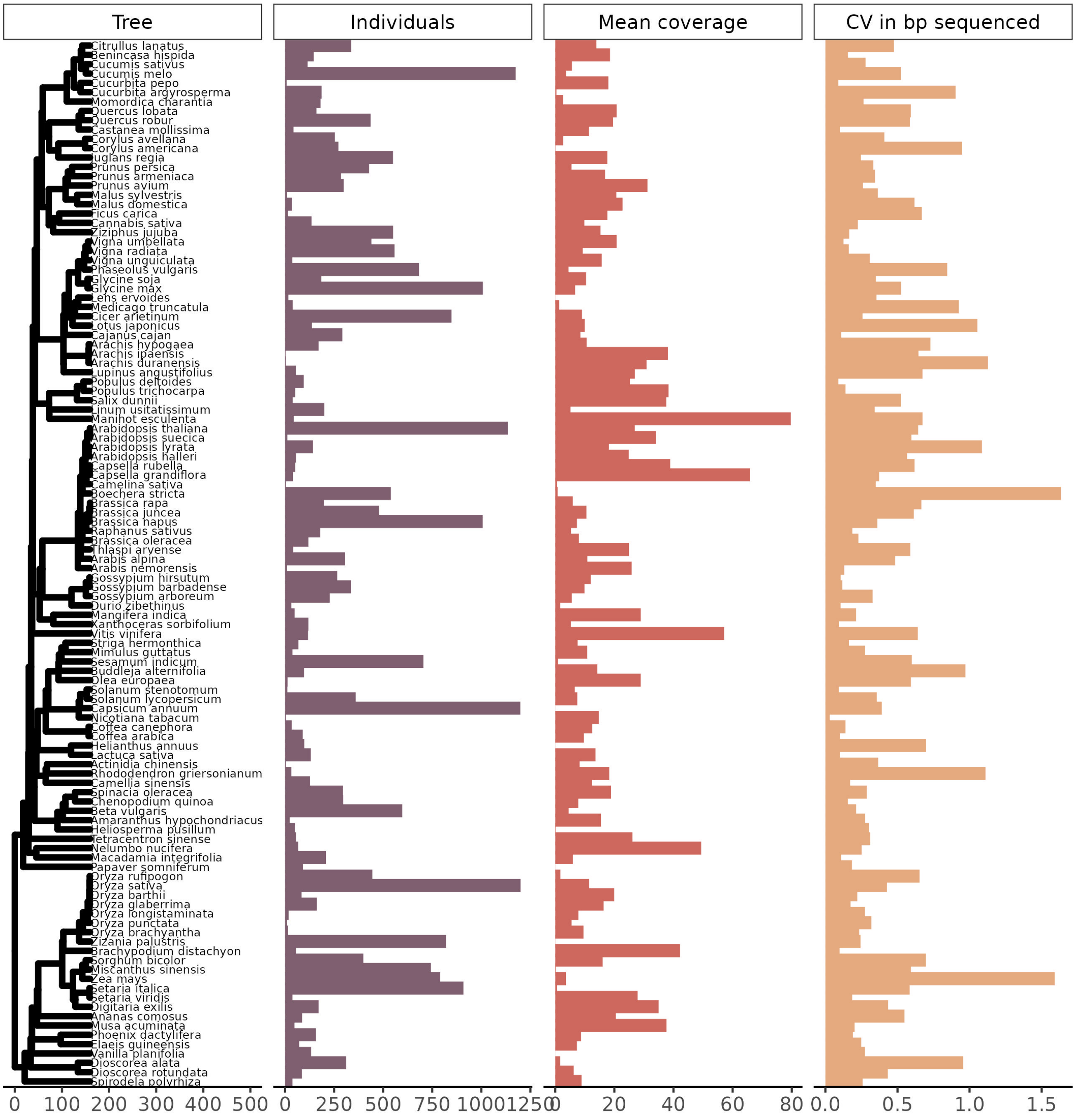


Figure S2: **Phylogenetic tree of species included in this study plotted alongside technical sequencing variables.** The phylogenetic tree scale is in millions of years. We limited the number of individuals sequenced per species to no more than 1200. CV stands for coefficient of variation and bp for base pairs.

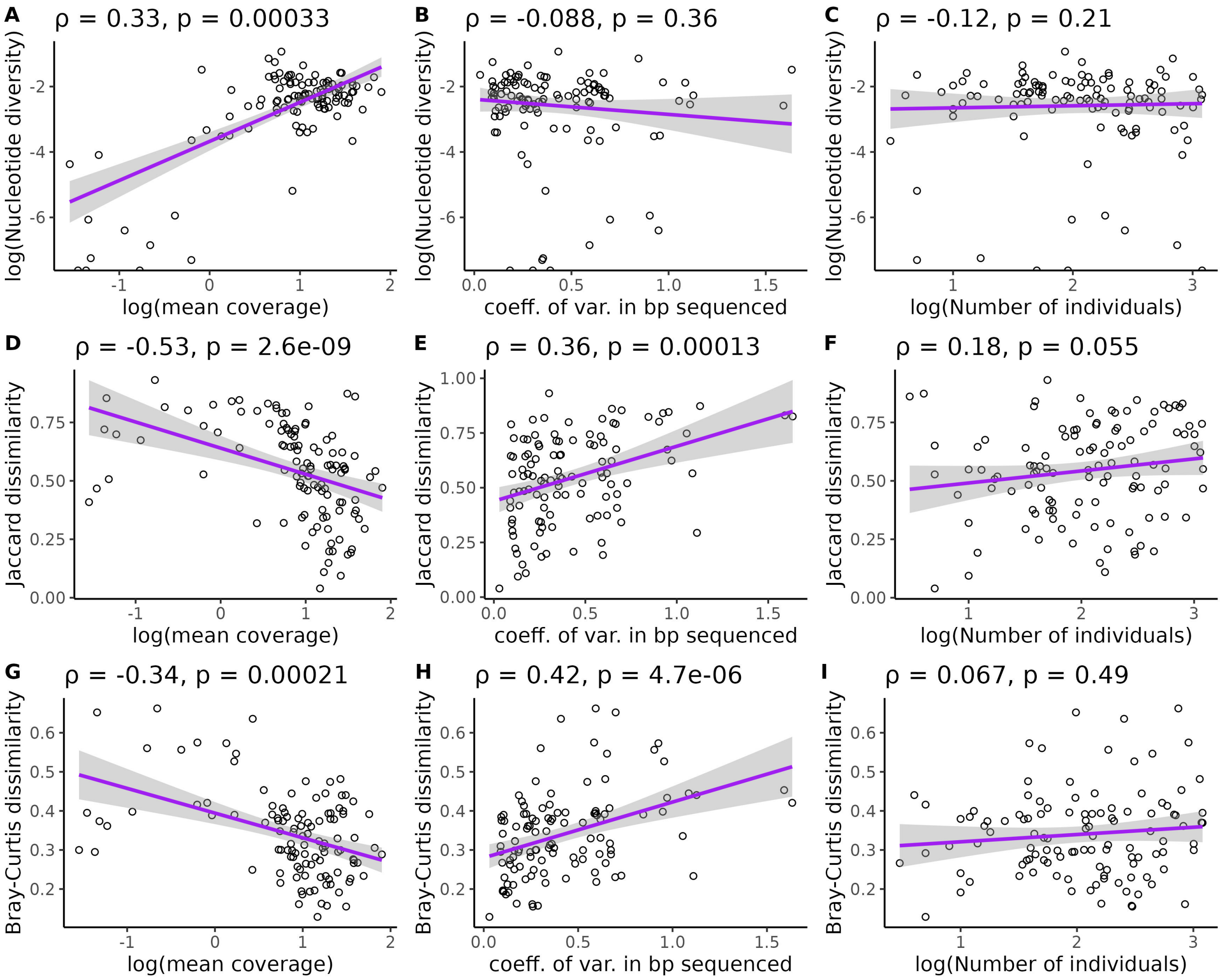


Figure S3: **Correlations between technical sequencing variables and diversity.** Each point is a species. Values across the top of each plot give the Spearman's correlation coefficient and p-value (testing whether the correlation differs from zero) for each pairwise relationship. Each pairwise plot includes one of three measures of diversity (Nucleotide diversity: A-C, Jaccard Dissimilarity: D-F, Bray-Curtis dissimilarity: G-I) and one of three technical sequencing variables (mean coverage: A, D, G; Coefficient of variation in bp sequenced: B, E, H; Number of individuals sequenced: C, F, I). Purple line is a loess smoothing line with 95% confidence intervals shaded in gray. All logarithms are base 10. Three species with nucleotide diversity values of 0 are omitted from plots involving nucleotide diversity.

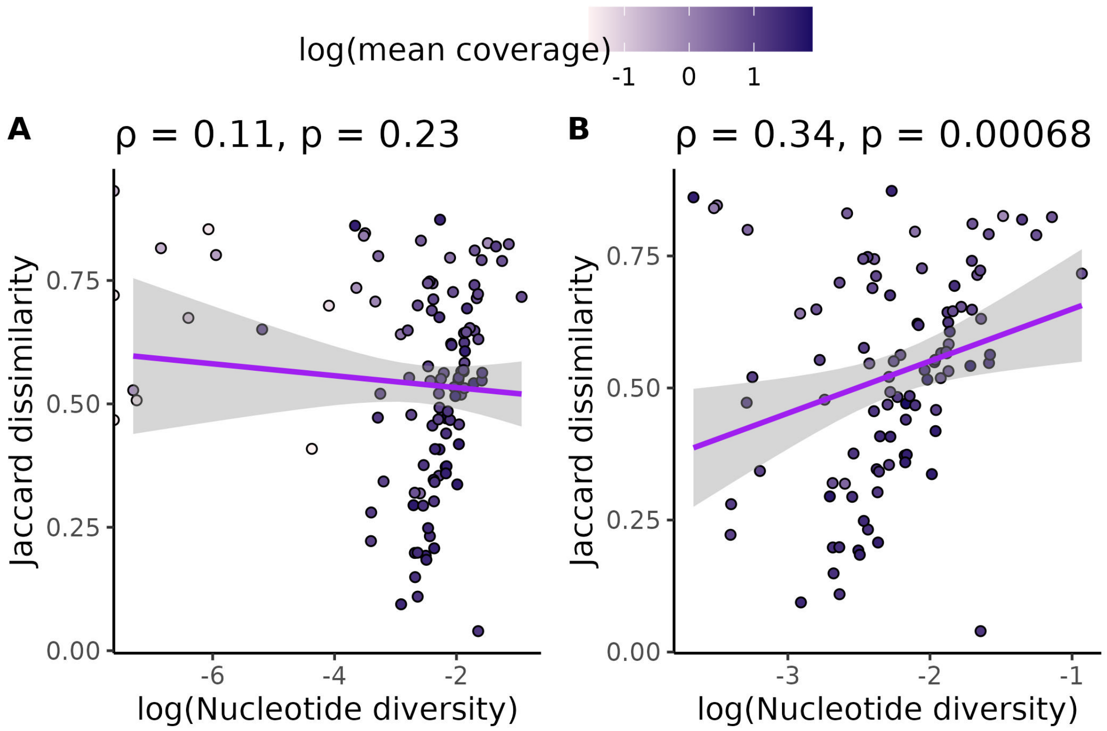


Figure S4: (A) shows the relationship between k-mer diversity (Jaccard dissimilarity) and nucleotide diversity without omitting species with $\leq$ 0.5x coverage or $\leq$ 1000 SNP calls. (B) shows the same relationship, except these species with low coverage or low numbers of SNP calls are omitted. Each data point is a species. All species' points are colored by the log of average genome-wide coverage per individual (base 10) for that species. Purple lines are loess smoothing curves with 95% confidence intervals shaded in gray. Values across the top of each plot are Spearman correlation coefficients ($\rho$) and p-values that test whether each correlation coefficient differs from zero. Three species with nucleotide diversity values of 0 are omitted from these plots.

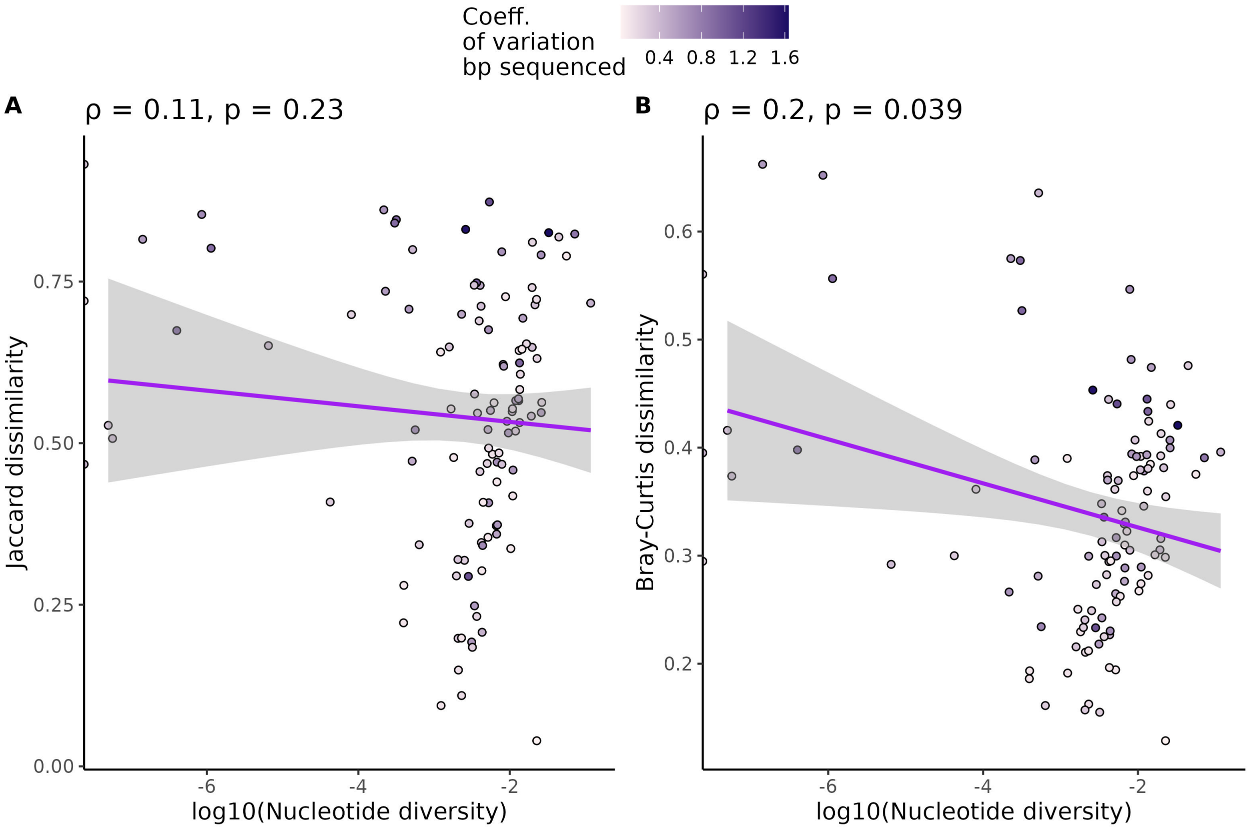


Figure S5: **Relationship between k-mer diversity, nucleotide diversity and variation in sequencing coverage.** Each point is a species. Values across the top of each plot give the Spearman's correlation coefficient and p-value (testing whether the correlation differs from zero) for each pairwise relationship. Purple line is a loess smoothing line with 95% confidence intervals shaded in gray. Points are colored by the coefficient of variation (mean/standard deviation) in bp sequenced for each species. Three species with nucleotide diversity values of 0 are omitted from the plots.

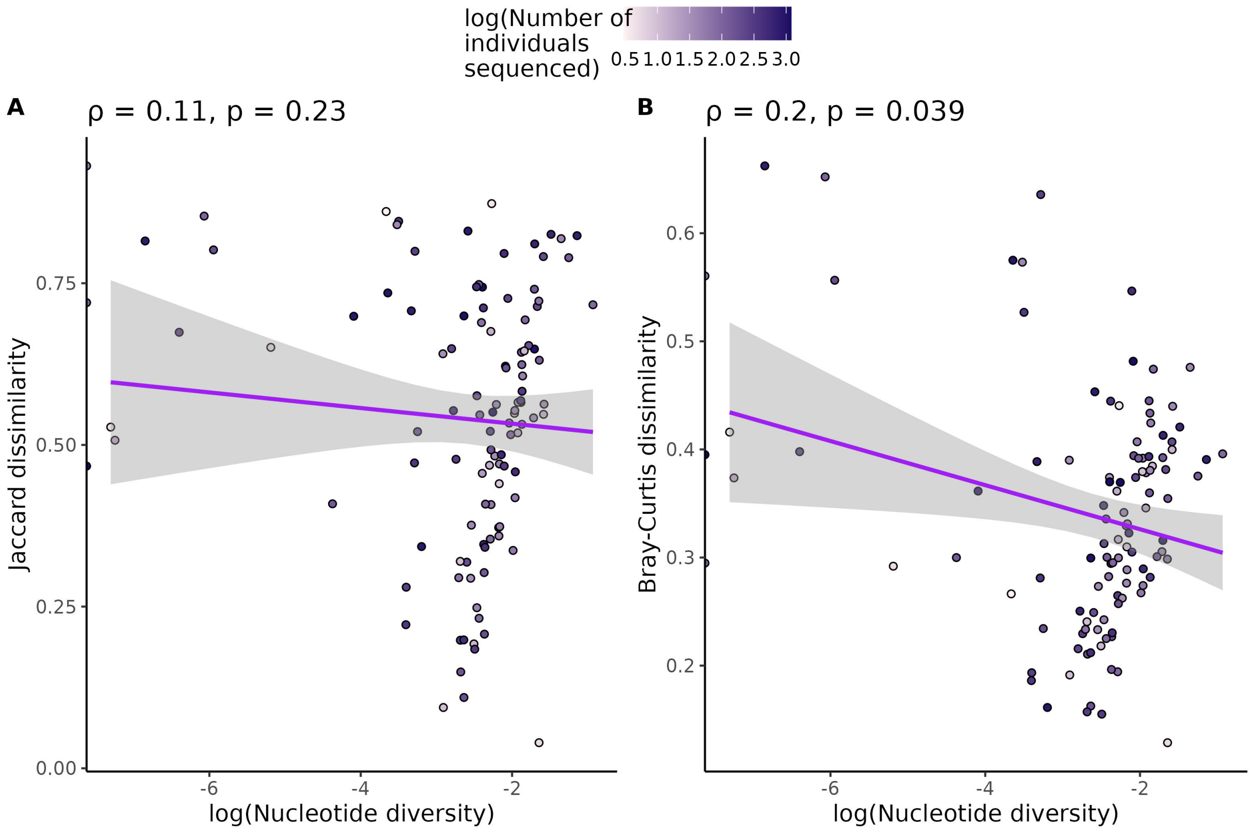


Figure S6: **Relationship between k-mer diversity, nucleotide diversity, and number of individuals sequenced.** Each point is a species. Values across the top of each plot give the Spearman's correlation coefficient and p-value (testing whether correlation differs from zero) for each pairwise relationship. Purple line is a loess smoothing line with 95% confidence intervals shaded in gray. Points are colored by the logarithm of the number of individuals sequenced within each species (base 10). Three species with nucleotide diversity values of 0 are omitted from the plots.

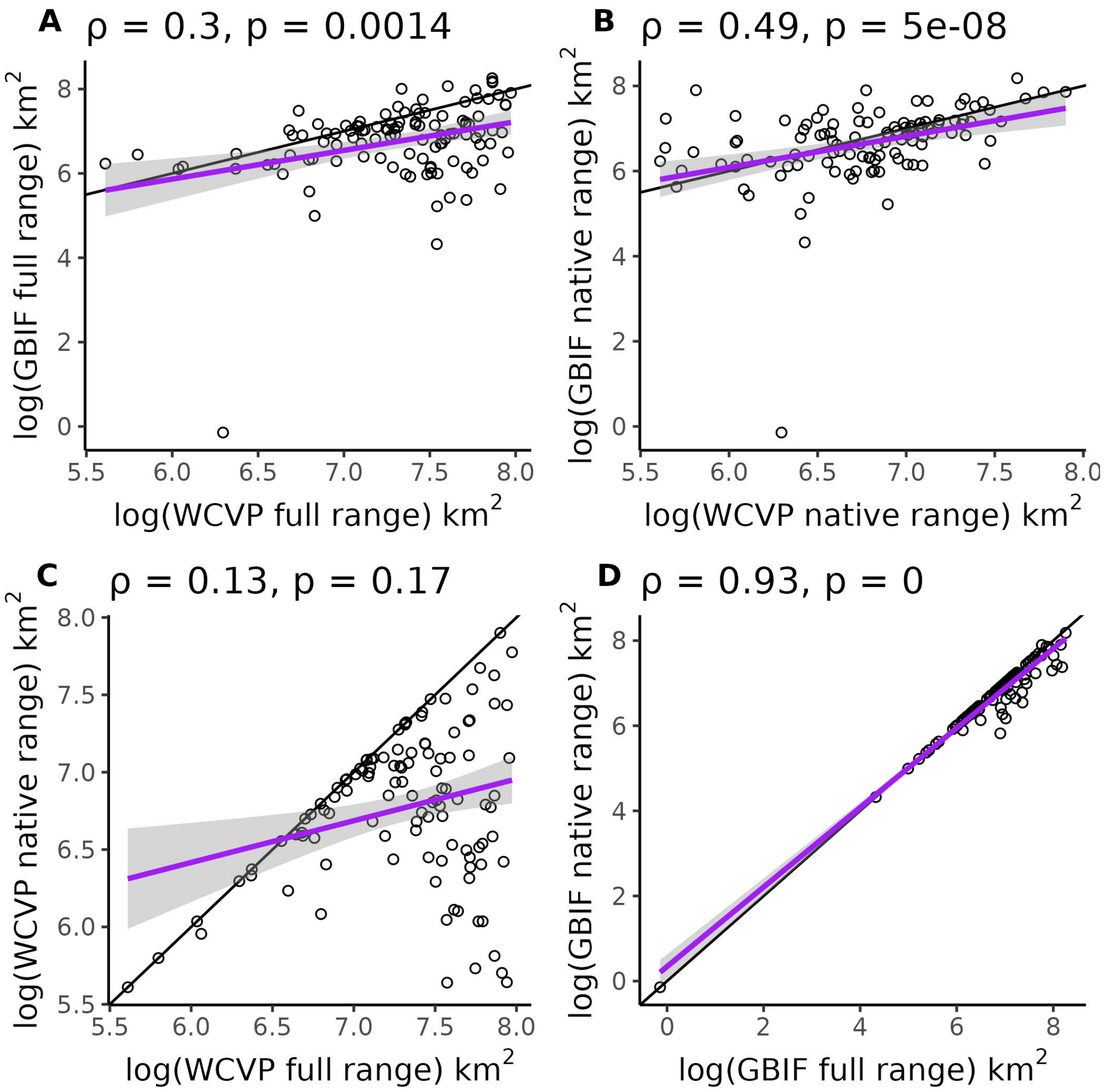


Figure S7: **Independent methods of range size estimation correlate with each other.** (A) Compares the size of total size of native and invaded ranges from GBIF occurrence data and WCVP range maps. (B) Compares native range estimates only (invaded ranges excluded) from GBIF occurrence data and WCVP range maps. (C) and (D) compare the full and native range estimates from WCVP range maps and GBIF occurrence data, respectively.Every data point is a species. The solid black lines are 1:1 reference lines where the different measures of range size are equal. The purple lines are linear regression lines with 95 % confidence intervals in grey shading. Values across the top of each plot give the Spearman's correlation coefficient ($\rho$) and p-value testing whether correlation differs from zero. One p-value is reported as zero because it was $< 2.2 \times 10^{-16}$, which is the limit of precision for doubles in R. All logarithms are base 10.

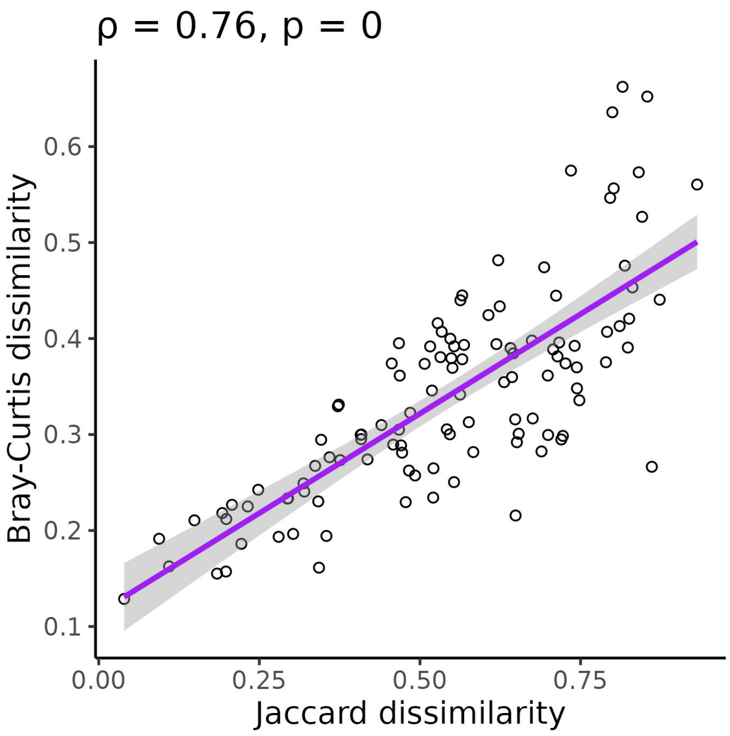


Figure S8: **Relationship between Bray-Curtis and Jaccard dissimilarity.** Each point is a species. Values across the top of each plot give the Spearman's correlation coefficient and p-value (testing whether correlation differs from zero). The p-value is reported as zero because it was $< 2.2 \times 10^{-16}$, which is the limit of precision for doubles in R. Line is a linear regression with 95% confidence intervals shaded in gray.

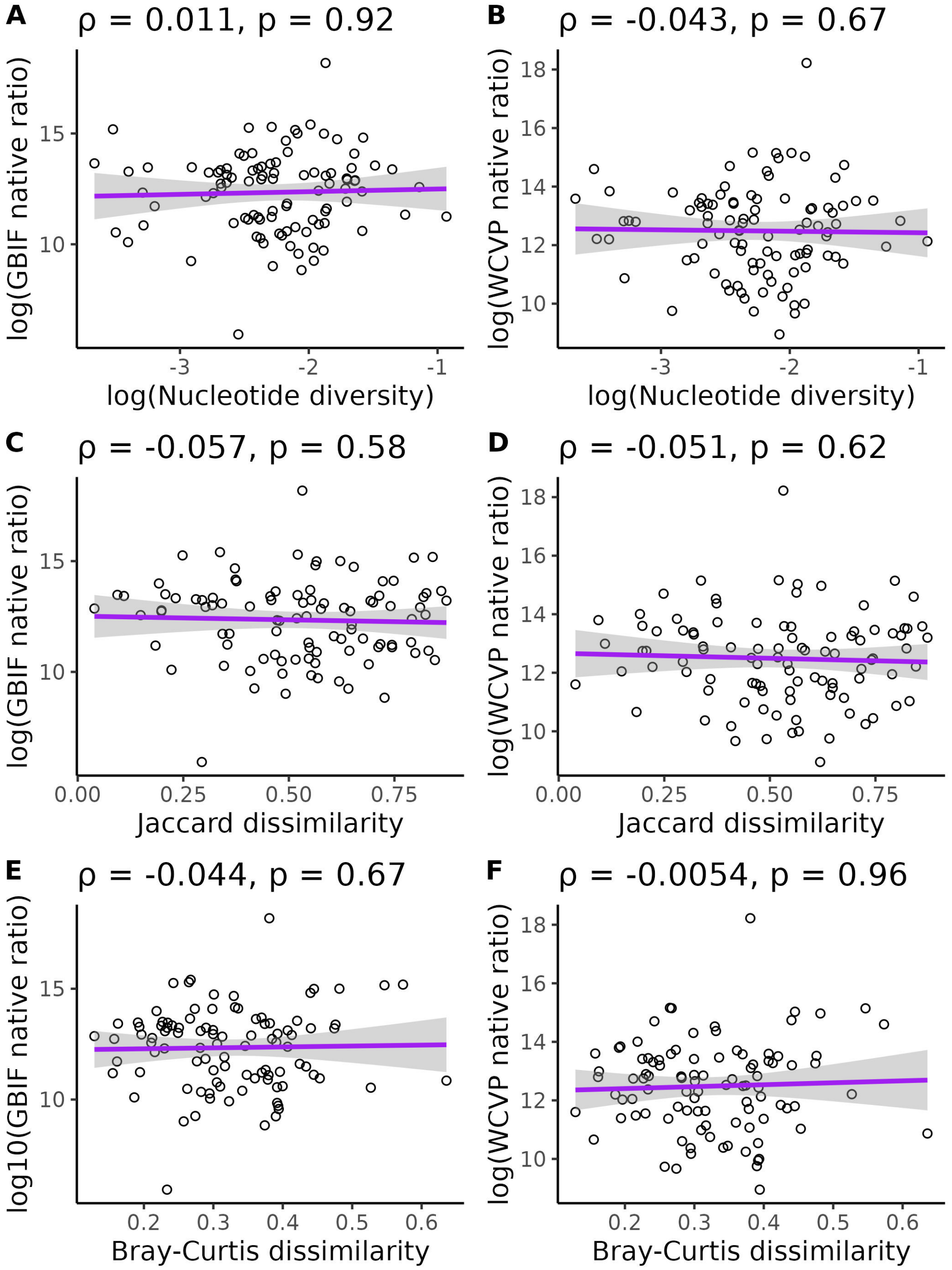


Figure S9: **Relationship between the range size-squared height ratio and diversity without correcting for evolutionary history, genome size, life cycle habit, mating system, or cultivation status.** Each point is a species. Values across the top of each plot give the Spearman's correlation coefficient and p-value (testing whether correlation differs from zero) for each pairwise relationship. Each pairwise relationship involves the native range size-squared height ratio, where native range size was estimated from GBIF occurrences (A, C, E), or WCVP range maps (B, D, F), and one of three diversity measures (Nucleotide diversity: A-B, Jaccard Dissimiarity: C-D, Bray-Curtis dissimilarity: E-F). Purple lines are loess smoothing lines with 95% confidence intervals shaded in gray. Three species with nucleotide diversity values of 0 are omitted from plots involving nucleotide diversity. All logarithms are base 10.

### 3 Supplemental figures: Population size proxy vs diversity relationships after controlling for phylogeny and life-history variables, but not controlling for genome size

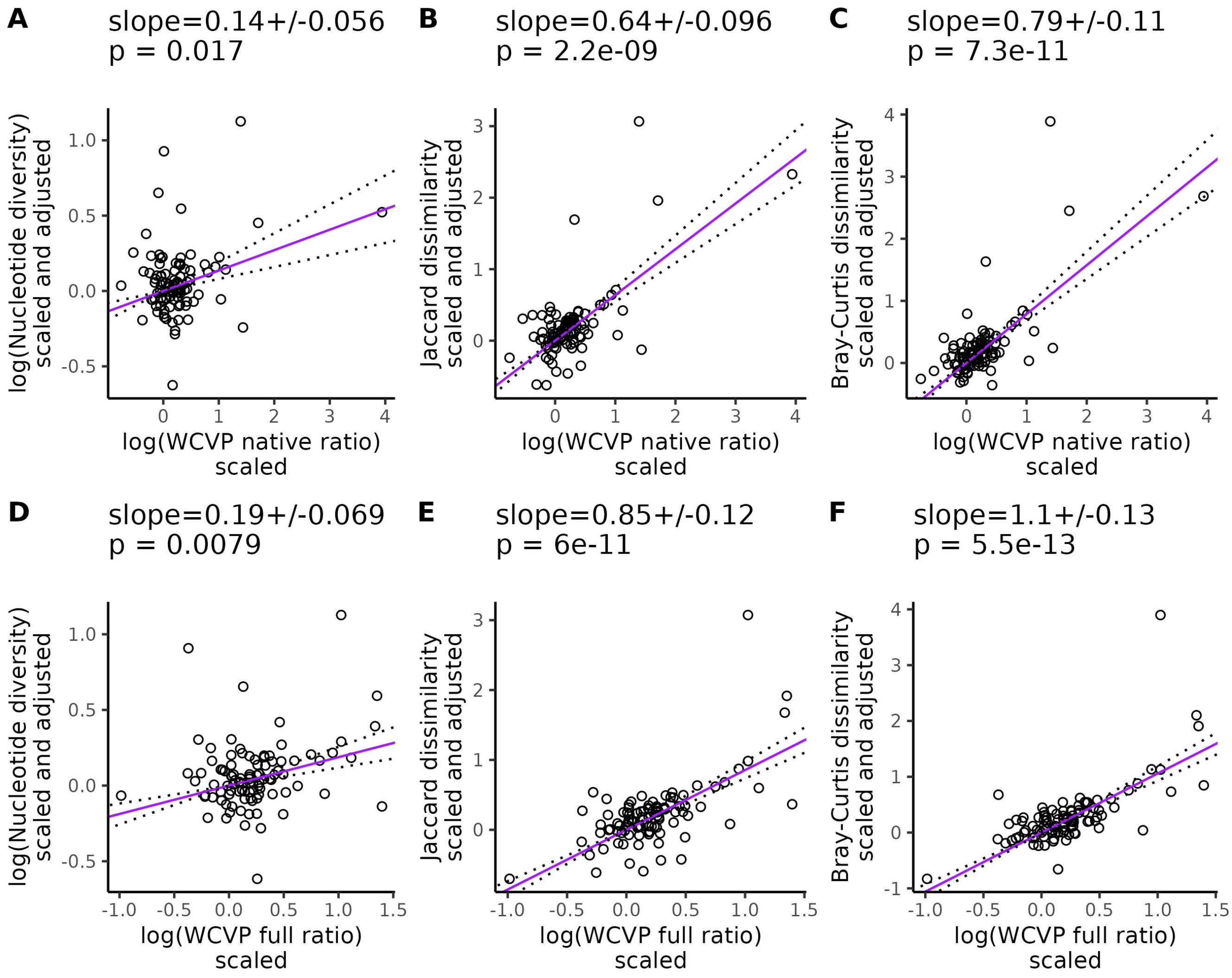


Figure S10: **Partial phylogenetic regression between WCVP population size proxy and diversity.** Each point is a species and only species with > 0.5x mean coverage and > 1000 variant sites were included in the regression. The regressions are organized according to whether invaded ranges were excluded (A-C) or included (D-F) in the range size-squared height ratio. Range size was estimated from WCVP range maps. Lines give the relationship between the pairs of plotted variables after controlling for mating system, life cycle habit, cultivation status, and evolutionary history. Before fitting the line, each response variable was scaled to a standard normal distribution (mean = 0, variance = 1), then multiplied by the inverse of the Cholesky decomposition of the phylogenetic variance-covariance matrix to correct for phylogenetic relationships. The values at the top of each plot give the slope of the partial regression ± one standard error and p-values testing whether the slopes differ from zero. Dotted lines show the partial regression slope ± one standard error. All logarithms are base 10.

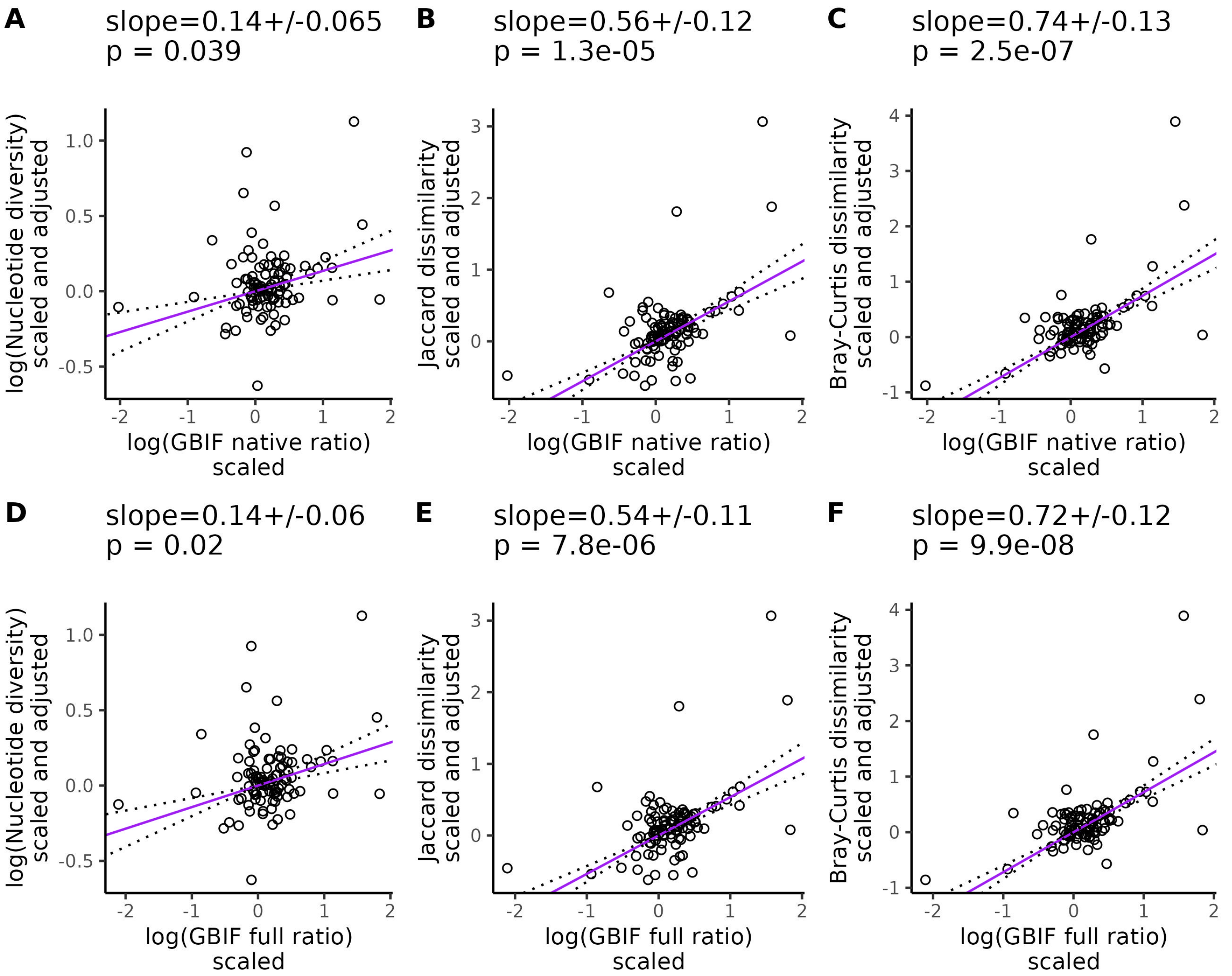


Figure S11: **Partial phylogenetic regression between GBIF population size proxy and diversity.** Each point is a species and only species with > 0.5x mean coverage and > 1000 variant sites were included in the regression. The regressions are organized according to whether invaded ranges were excluded (A-C) or included (D-F) in the range size-squared height ratio. Range size was estimated from GBIF occurrence data. Lines give the relationship between the pairs of plotted variables after controlling for mating system, life cycle habit, cultivation status, and evolutionary history. Before fitting the line, each response variable was scaled to a standard normal distribution (mean = 0, variance = 1), then multiplied by the inverse of the Cholesky decomposition of the phylogenetic variance-covariance matrix to correct for phylogenetic relationships. The values at the top of each plot give the slope of the partial regression ± one standard error and p-values testing whether the slopes differ from zero. Dotted lines show the partial regression slope ± one standard error. All logarithms are base 10.

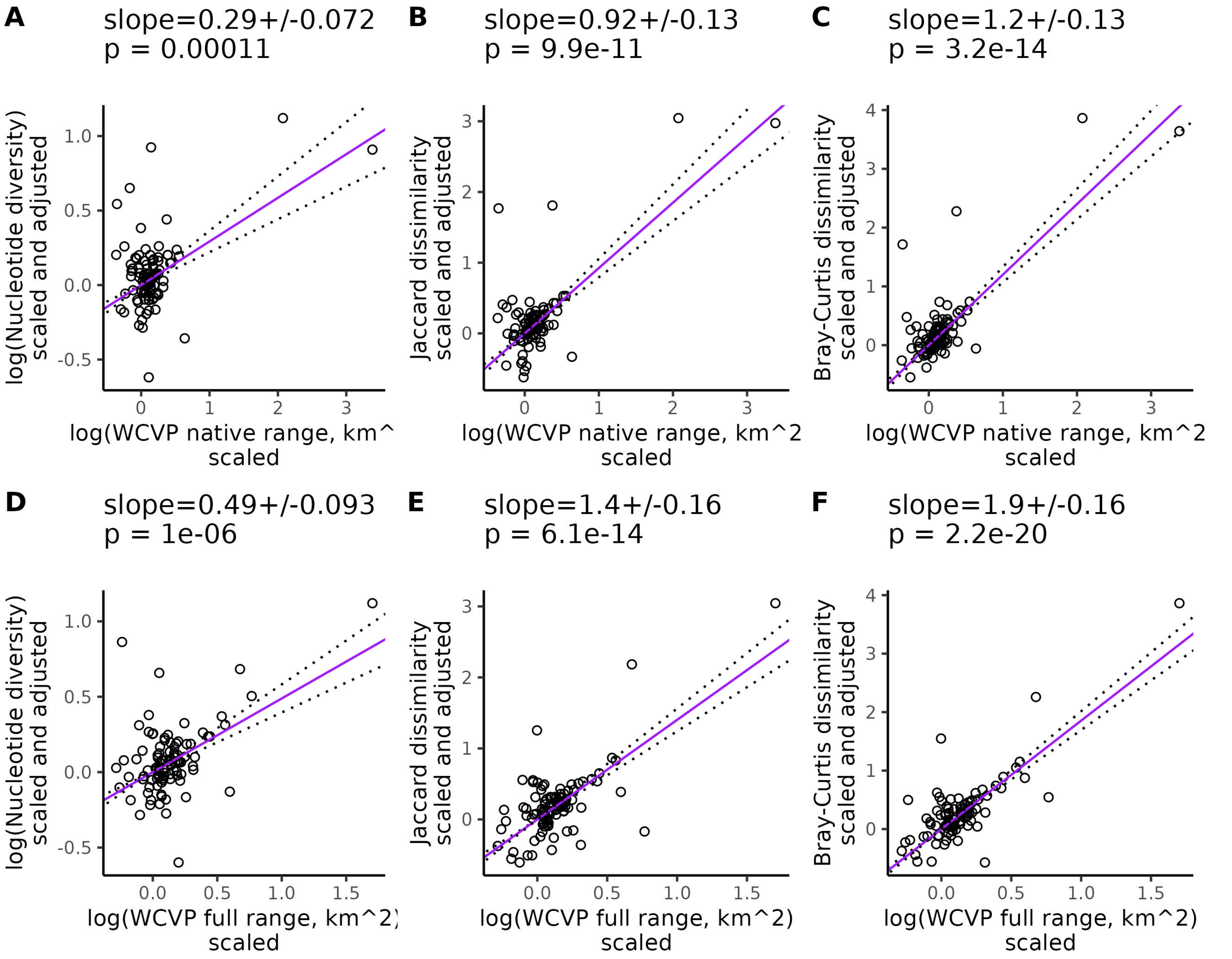


Figure S12: **Partial phylogenetic regression between WCVP range size and diversity.** Each point is a species and only species with > 0.5x mean coverage and > 1000 variant sites were included in the regression. The regressions are organized according to whether invaded ranges were excluded (A-C) or included (D-F) in the range size estimates. Range size was estimated from WCVP range maps. Lines give the relationship between the pairs of plotted variables after controlling for mating system, life cycle habit, cultivation status, and evolutionary history. Before fitting the line, each response variable was scaled to a standard normal distribution (mean = 0, variance = 1), then multiplied by the inverse of the Cholesky decomposition of the phylogenetic variance-covariance matrix to correct for phylogenetic relationships. The values at the top of each plot give the slope of the partial regression ± one standard error and p-values testing whether the slopes differ from zero. Dotted lines show the partial regression slope ± one standard error. All logarithms are base 10.

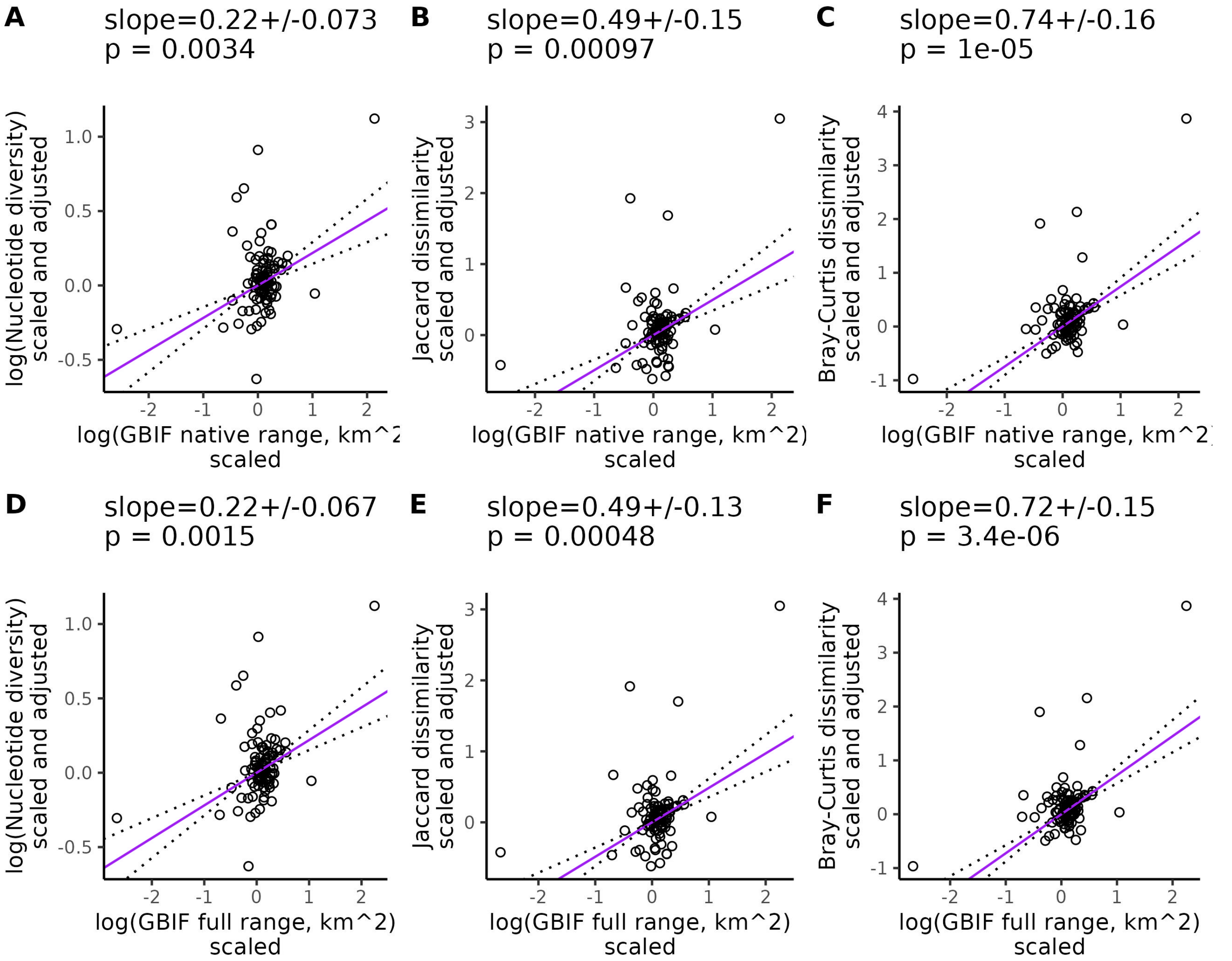


Figure S13: **Partial phylogenetic regression between GBIF range size and diversity.** Each point is a species and only species with > 0.5x mean coverage and > 1000 variant sites were included in the regression. The regressions are organized according to whether invaded ranges were excluded (A-C) or included (D-F) in the range size estimates. Range size was estimated from GBIF occurrence data. Lines give the relationship between the pairs of plotted variables after controlling for mating system, life cycle habit, cultivation status, and evolutionary history. Before fitting the line, each response variable was scaled to a standard normal distribution (mean = 0, variance = 1), then multiplied by the inverse of the Cholesky decomposition of the phylogenetic variance-covariance matrix to correct for phylogenetic relationships. The values at the top of each plot give the slope of the partial regression ± one standard error and p-values testing whether the slopes differ from zero. Dotted lines show the partial regression slope ± one standard error. All logarithms are base 10.

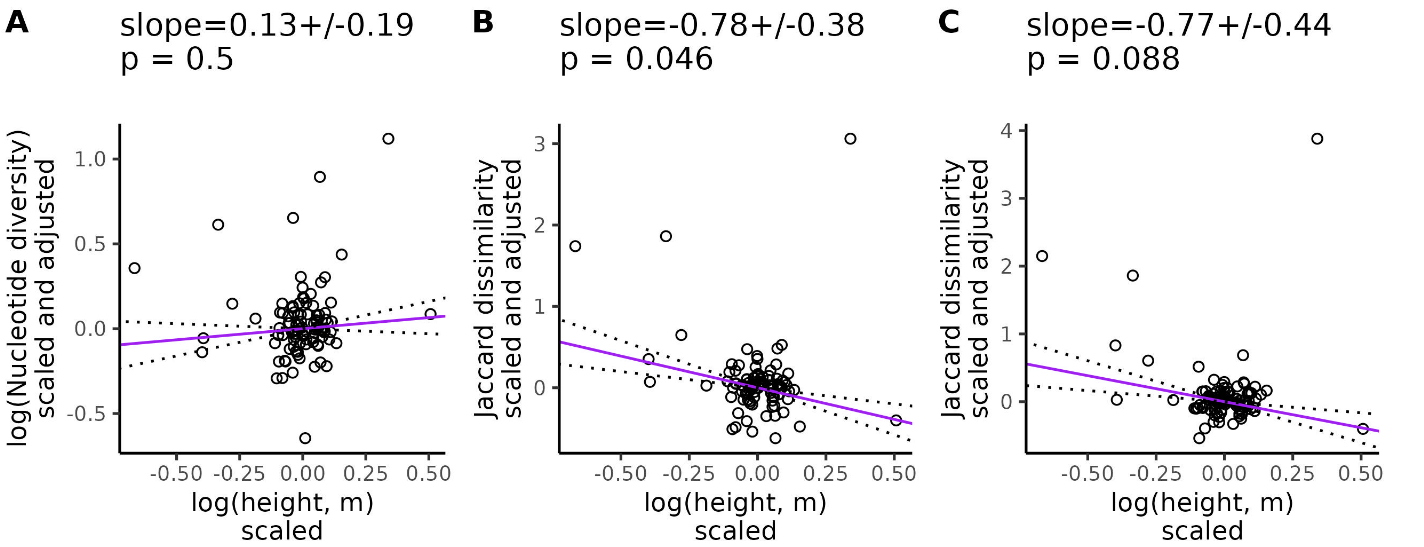


Figure S14: **Partial phylogenetic regression between height and diversity.** Each point is a species and only species with > 0.5x mean coverage and > 1000 variant sites were included in the regression. Lines give the relationship between the pairs of plotted variables after controlling for mating system, life cycle habit, cultivation status, and evolutionary history. The statistics across the top of each plot give the value of the slope of the lines (± the standard error), the p-value testing whether the slope differs from zero. Before fitting the line, each response variable was scaled to a standard normal distribution (mean = 0, variance = 1), then multiplied by the inverse of the Cholesky decomposition of the phylogenetic variance-covariance matrix to correct for phylogenetic relationships. All logarithms are base 10.

### 4 Supplemental figures: Population size proxy vs diversity relationships after controlling for phylogeny, life-history variables, and genome size

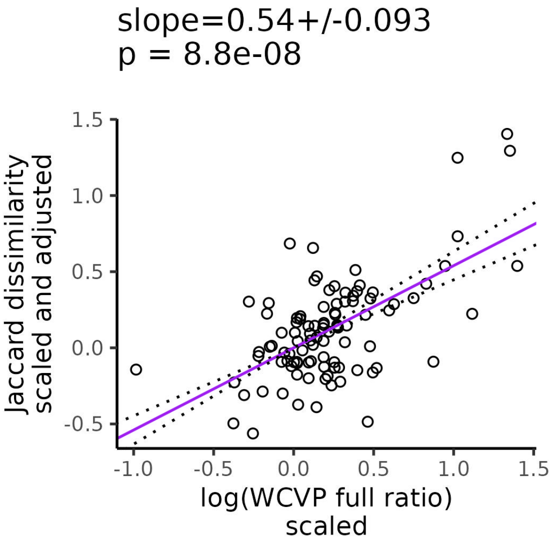


Figure S15: **Partial phylogenetic regression between WCVP full range size-squared height ratio and Jaccard dissimilarity, controlling for genome size.** Each point is a species and only species with $> 0.5$x mean coverage and $> 1000$ variant sites were included in the regression. WCVP full ratio gives the ratio of range size to squared plant height, where range size includes invaded ranges and is estimated from WCVP range maps. The partial regression controls for genome size, mating system, life cycle habit, cultivation status, and evolutionary history. Before fitting the line, each response variable was scaled to a standard normal distribution (mean = 0, variance = 1), then multiplied by the inverse of the Cholesky decomposition of the phylogenetic variance-covariance matrix to correct for phylogenetic relationships. The values at the top of the plot give the slope of the partial regression $\pm$ one standard error and p-values testing whether the slope differ from zero. Dotted lines show the partial regression slope $\pm$ one standard error. All logarithms are base 10.

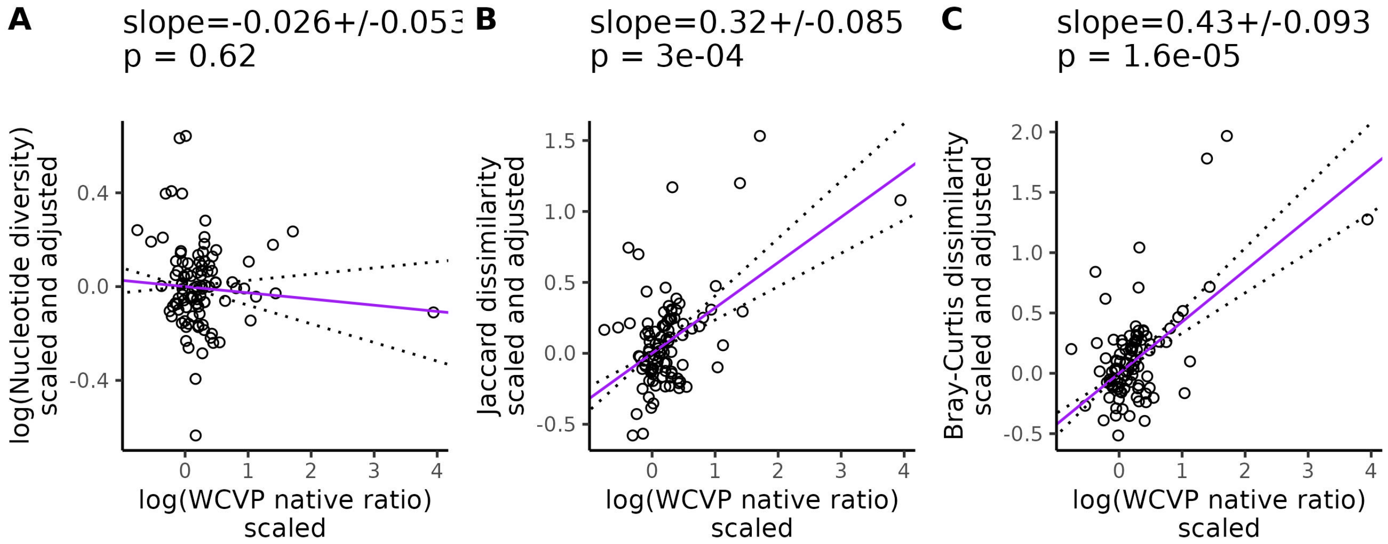


Figure S16: **Partial phylogenetic regression between WCVP native range size-squared height ratio and diversity, controlling for genome size.** Each point is a species and only species with > 0.5x mean coverage and > 1000 variant sites were included in the regression. WCVP native ratio gives the ratio of range size to squared plant height, where range size excludes invaded ranges and is estimated from WCVP range maps. The partial regression controls for genome size, mating system, life cycle habit, cultivation status, and evolutionary history. Before fitting the line, each response variable was scaled to a standard normal distribution (mean = 0, variance = 1), then multiplied by the inverse of the Cholesky decomposition of the phylogenetic variance-covariance matrix to correct for phylogenetic relationships. The values at the top of the plot give the slope of the partial regression ± one standard error and p-values testing whether the slope differ from zero. Dotted lines show the partial regression slope ± one standard error. All logarithms are base 10.

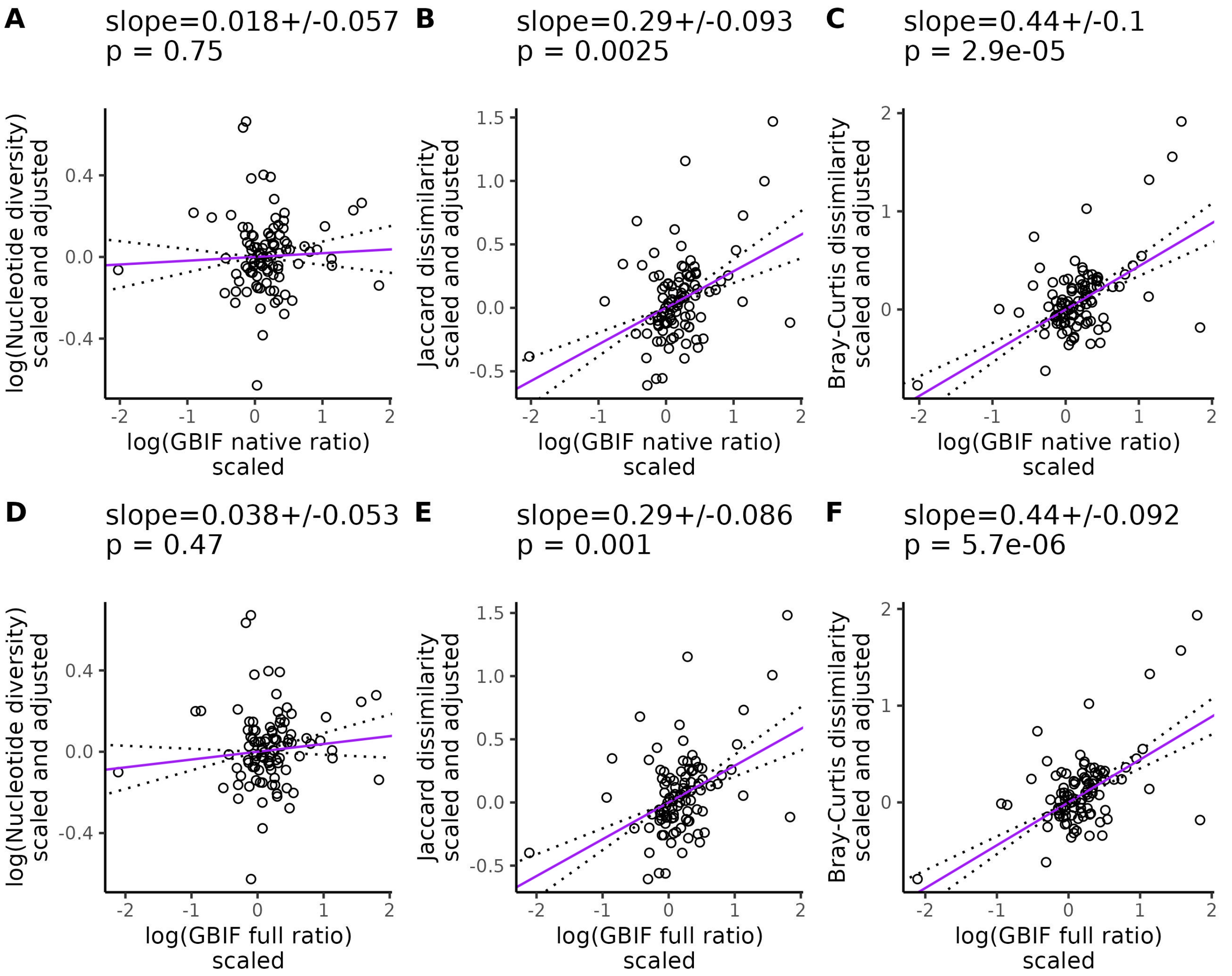


Figure S17: **Partial phylogenetic regression between GBIF range size-squared height ratio and diversity, controlling for genome size.** Each point is a species and only species with $>$ 0.5x mean coverage and $>$ 1000 variant sites were included in the regression. The regressions are organized according to whether invaded ranges were excluded (A-C) or included (D-F) in the range size-squared height ratio. Lines give the relationship between the pairs of plotted variables after controlling for mating system, life cycle habit, cultivation status, genome size, and evolutionary history. The statistics across the top of each plot give the value of the slope of the lines ($\pm$ the standard error) and the p-value testing whether the slope differs from zero. Before fitting the line, each response variable was scaled to a standard normal distribution (mean = 0, variance = 1), then multiplied by the inverse of the Cholesky decomposition of the phylogenetic variance-covariance matrix to correct for phylogenetic relationships. Dotted lines show the partial regression slope $\pm$ one standard error. All logarithms are base 10.

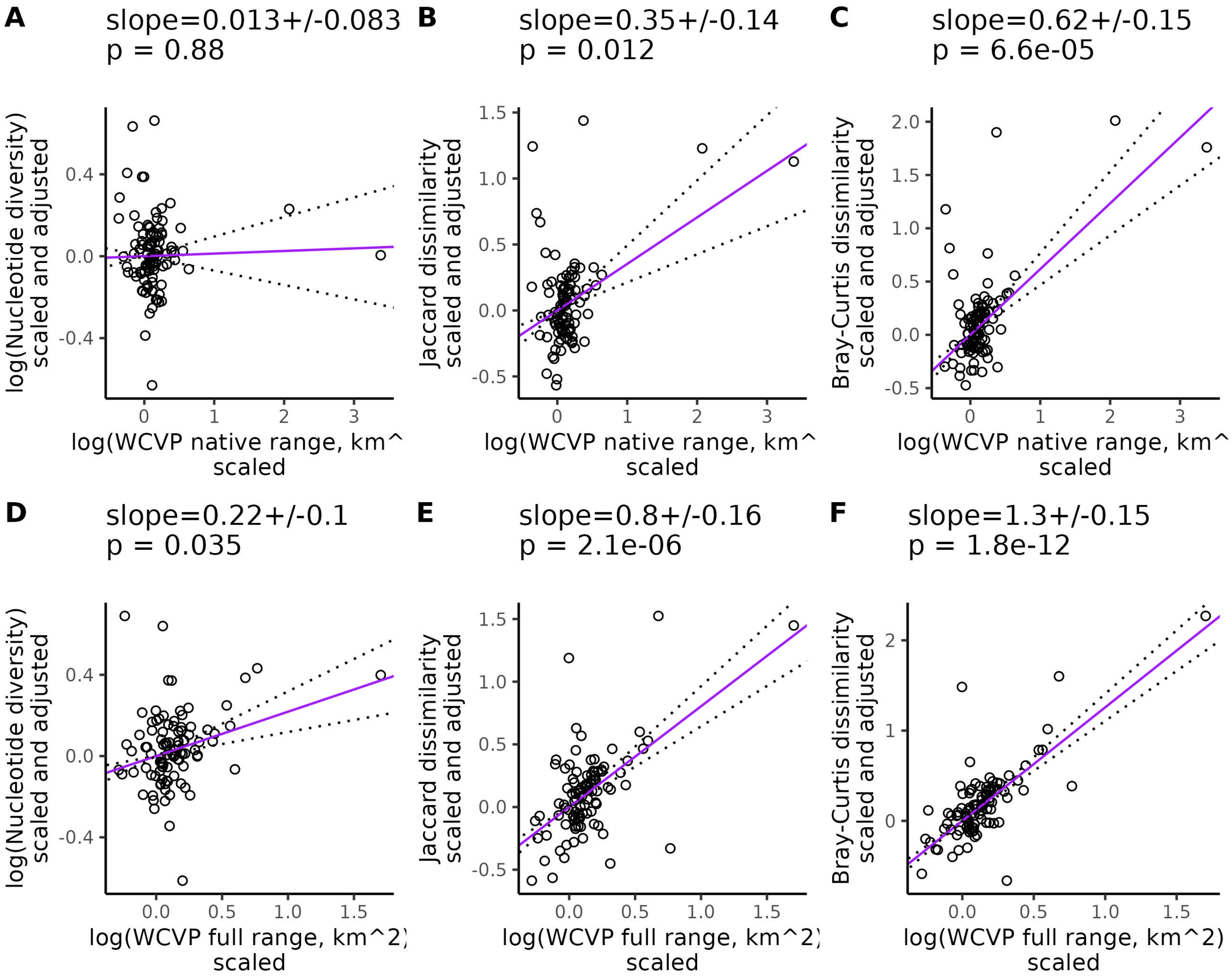


Figure S18: **Partial phylogenetic regression between WCVP range size and diversity, controlling for genome size.** Each point is a species and only species with > 0.5x mean coverage and > 1000 variant sites were included in the regression. The regressions are organized according to whether invaded ranges were excluded (A-C) or included (D-F) in the range size estimates. Lines give the relationship between the pairs of plotted variables after controlling for mating system, life cycle habit, cultivation status, genome size, and evolutionary history. The statistics across the top of each plot give the value of the slope of the lines (± the standard error) and the p-value testing whether the slope differs from zero. Before fitting the line, each response variable was scaled to a standard normal distribution (mean = 0, variance = 1), then multiplied by the inverse of the Cholesky decomposition of the phylogenetic variance-covariance matrix to correct for phylogenetic relationships. Dotted lines show the partial regression slope ± one standard error. All logarithms are base 10.

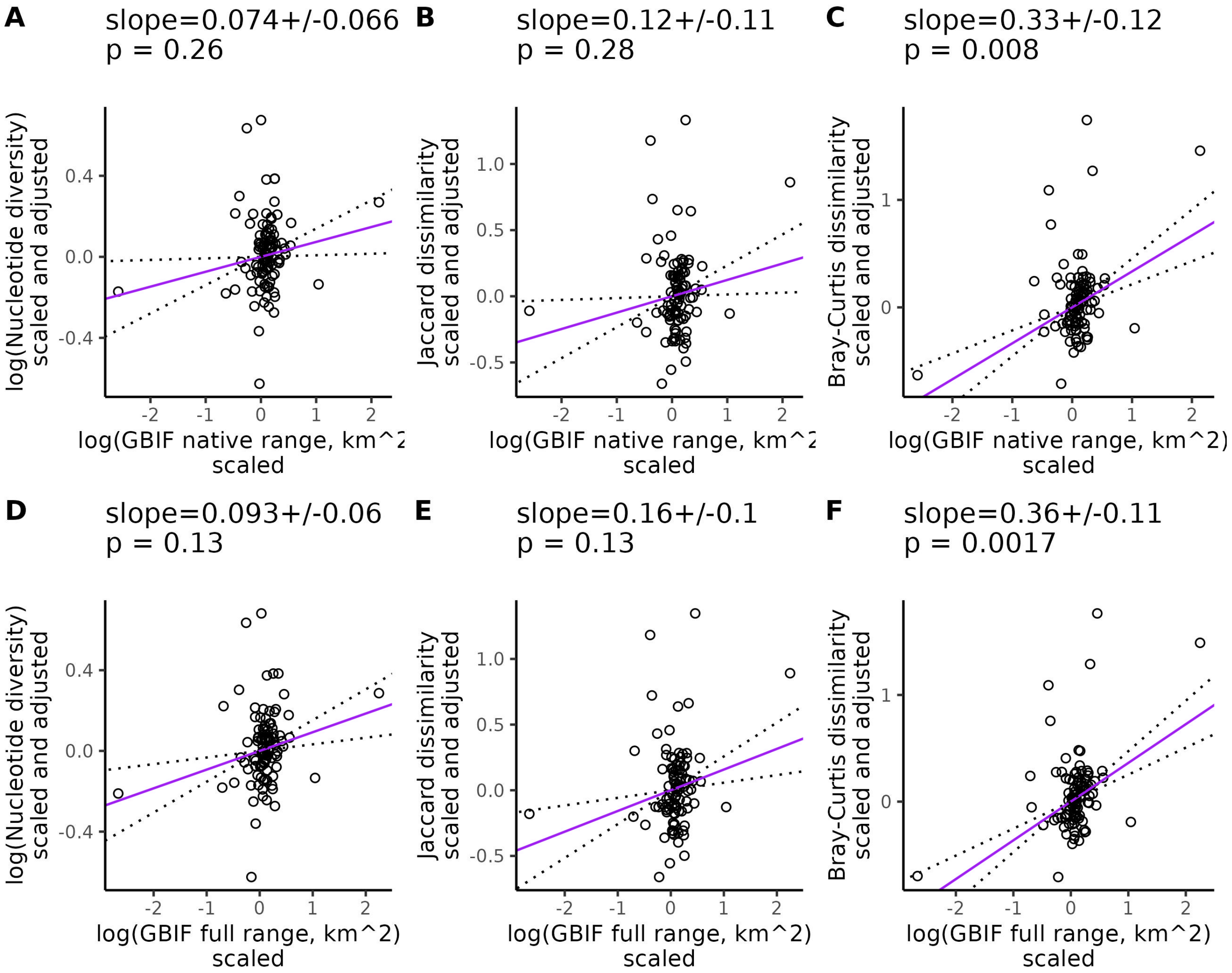


Figure S19: **Partial phylogenetic regression between GBIF range size and diversity, controlling for genome size.** Each point is a species and only species with $> 0.5$x mean coverage and $> 1000$ variant sites were included in the regression. The regressions are organized according to whether invaded ranges were excluded (A-C) or included (D-F) in the range size estimates. Lines give the relationship between the pairs of plotted variables after controlling for mating system, life cycle habit, cultivation status, genome size, and evolutionary history. The statistics across the top of each plot give the value of the slope of the lines ($\pm$ the standard error) and the p-value testing whether the slope differs from zero. Before fitting the line, each response variable was scaled to a standard normal distribution (mean = 0, variance = 1), then multiplied by the inverse of the Cholesky decomposition of the phylogenetic variance-covariance matrix to correct for phylogenetic relationships. Dotted lines show the partial regression slope $\pm$ one standard error. All logarithms are base 10.

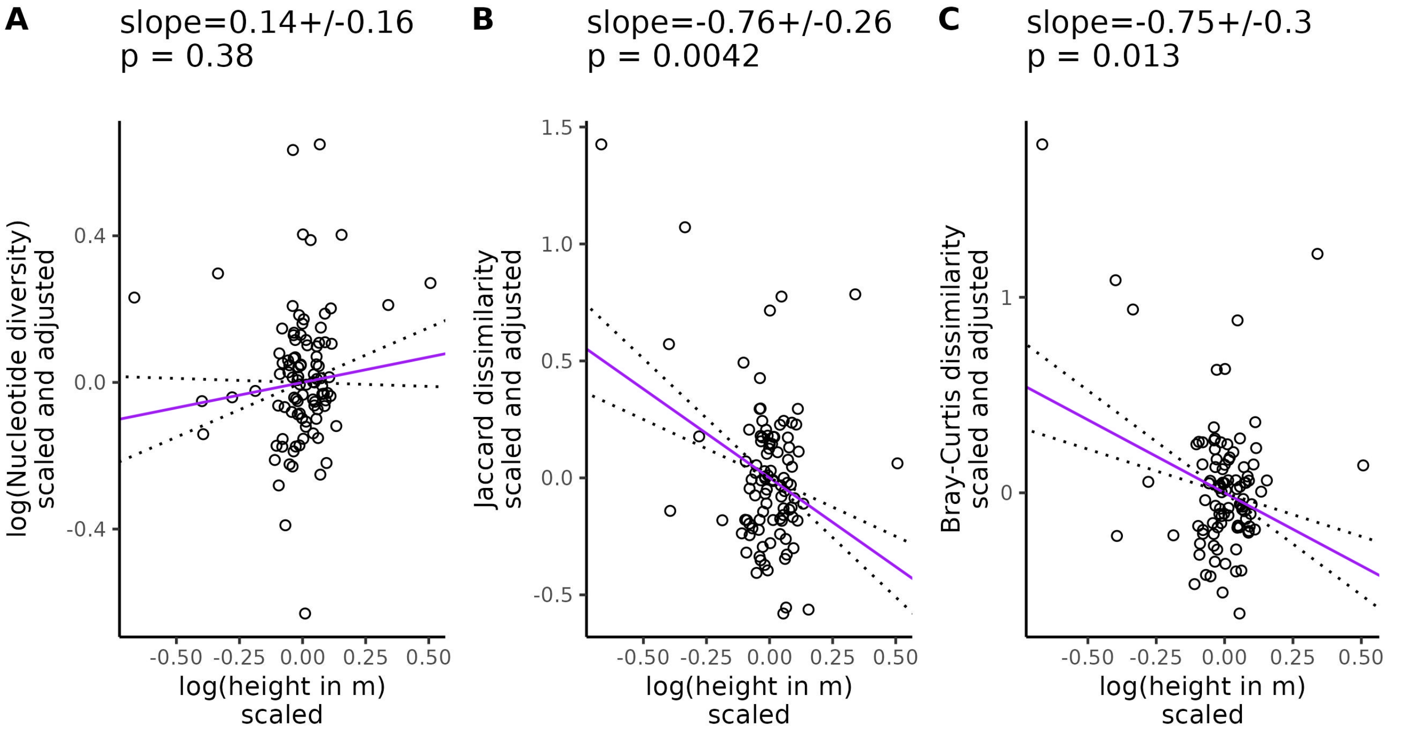


Figure S20: **Partial phylogenetic regression between height and diversity, controlling for genome size.** Each point is a species and only species with > 0.5x mean coverage and > 1000 variant sites were included in the regression. Lines give the relationship between the pairs of plotted variables after controlling for mating system, life cycle habit, cultivation status, genome size, and evolutionary history. The statistics across the top of each plot give the value of the slope of the lines ($\pm$ the standard error) and the p-value testing whether the slope differs from zero. Before fitting the line, each response variable was scaled to a standard normal distribution (mean = 0, variance = 1), then multiplied by the inverse of the Cholesky decomposition of the phylogenetic variance-covariance matrix to correct for phylogenetic relationships. Dotted lines show the partial regression slope $\pm$ one standard error. All logarithms are base 10.

### 5 Supplemental figures: Genome size vs diversity relationships after controlling for population size proxies, phylogeny, and life-history variables

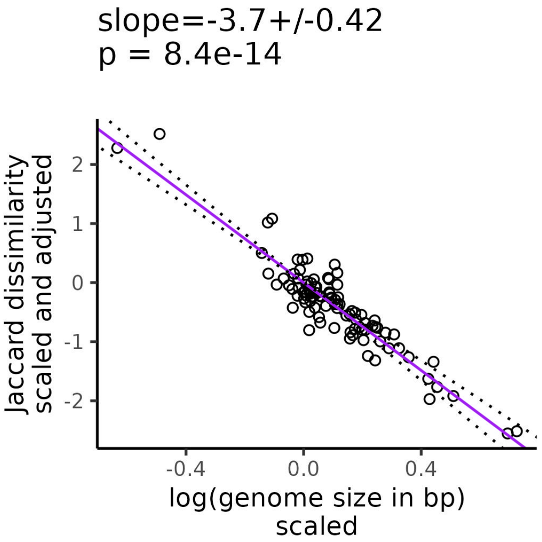


Figure S21: **Partial phylogenetic regression between Jaccard dissimilarity and genome size, controlling for WCVP full range size-height ratio.** Each point is a species and only species with > 0.5x mean coverage and > 1000 variant sites were included in the regression. WCVP full ratio gives the ratio of range size to squared plant height, where range size includes invaded ranges and is estimated from WCVP range maps. The partial regression controls for range size-squared height ratio, mating system, life cycle habit, cultivation status, and evolutionary history. Before fitting the line, each response variable was scaled to a standard normal distribution (mean = 0, variance = 1), then multiplied by the inverse of the Cholesky decomposition of the phylogenetic variance-covariance matrix to correct for phylogenetic relationships. The values at the top of the plot give the slope of the partial regression ± one standard error and p-values testing whether the slope differ from zero. Dotted lines show the partial regression slope ± one standard error. All logarithms are base 10.

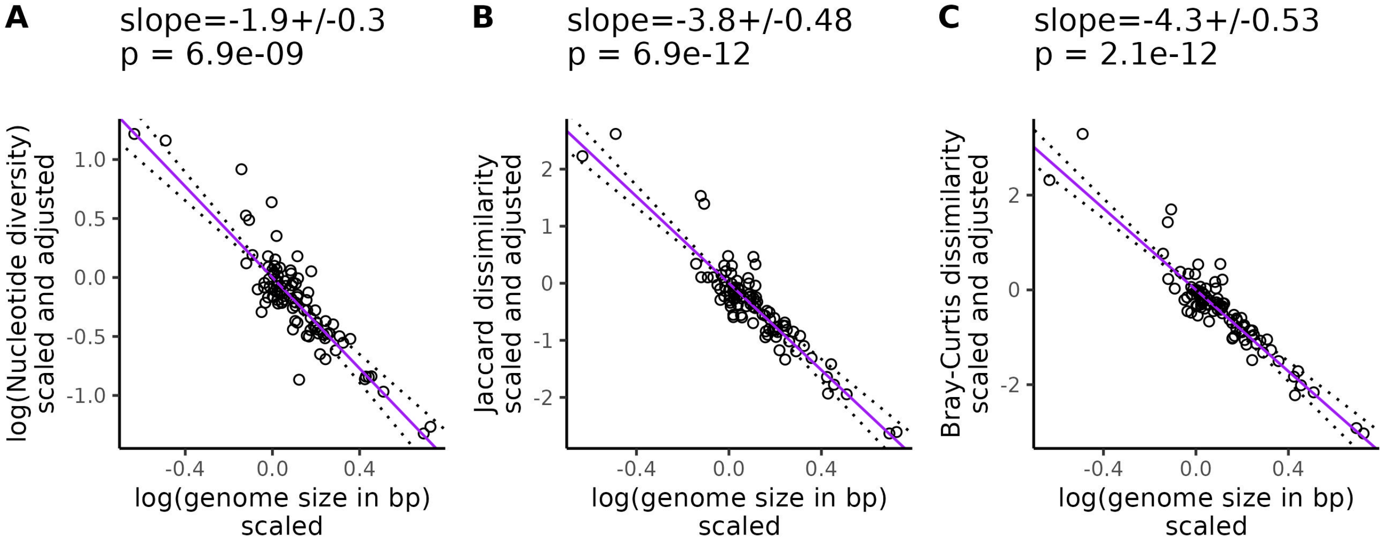


Figure S22: **Partial phylogenetic regression between diversity and genome size, controlling for WCVP native range size-height ratio.** Each point is a species and only species with $>$ 0.5x mean coverage and $>$ 1000 variant sites were included in the regression. WCVP native ratio gives the ratio of range size to squared plant height, where range size excludes invaded ranges and is estimated from WCVP range maps. The partial regression controls for genome size, mating system, life cycle habit, cultivation status, and evolutionary history. Before fitting the line, each response variable was scaled to a standard normal distribution (mean = 0, variance = 1), then multiplied by the inverse of the Cholesky decomposition of the phylogenetic variance-covariance matrix to correct for phylogenetic relationships. The values at the top of the plot give the slope of the partial regression $\pm$ one standard error and p-values testing whether the slope differ from zero. Dotted lines show the partial regression slope $\pm$ one standard error. All logarithms are base 10.

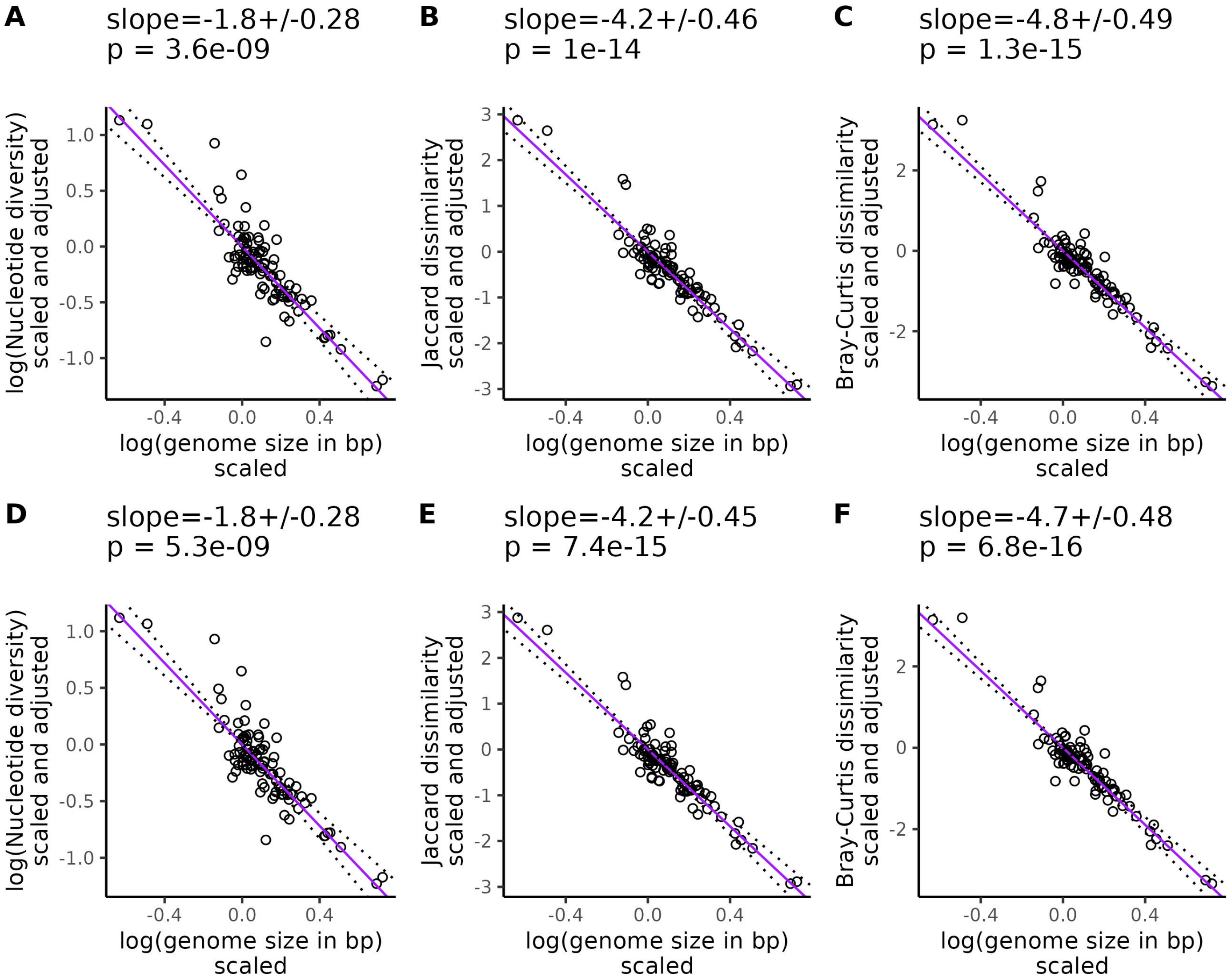


Figure S23: **Partial phylogenetic regression between diversity and genome size, controlling for GBIF range size-height ratio.** Each point is a species and only species with > 0.5x mean coverage and > 1000 variant sites were included in the regression. The regressions are organized according to whether invaded ranges were excluded (A-C) or included (D-F) in the range size-squared height ratio. Lines give the relationship between the pairs of plotted variables after controlling for mating system, life cycle habit, cultivation status, range size-squared height ratio, and evolutionary history. The statistics across the top of each plot give the value of the slope of the lines ($\pm$ the standard error) and the p-value testing whether the slope differs from zero. Before fitting the line, each response variable was scaled to a standard normal distribution (mean = 0, variance = 1), then multiplied by the inverse of the Cholesky decomposition of the phylogenetic variance-covariance matrix to correct for phylogenetic relationships. Dotted lines show the partial regression slope $\pm$ one standard error. All logarithms are base 10.

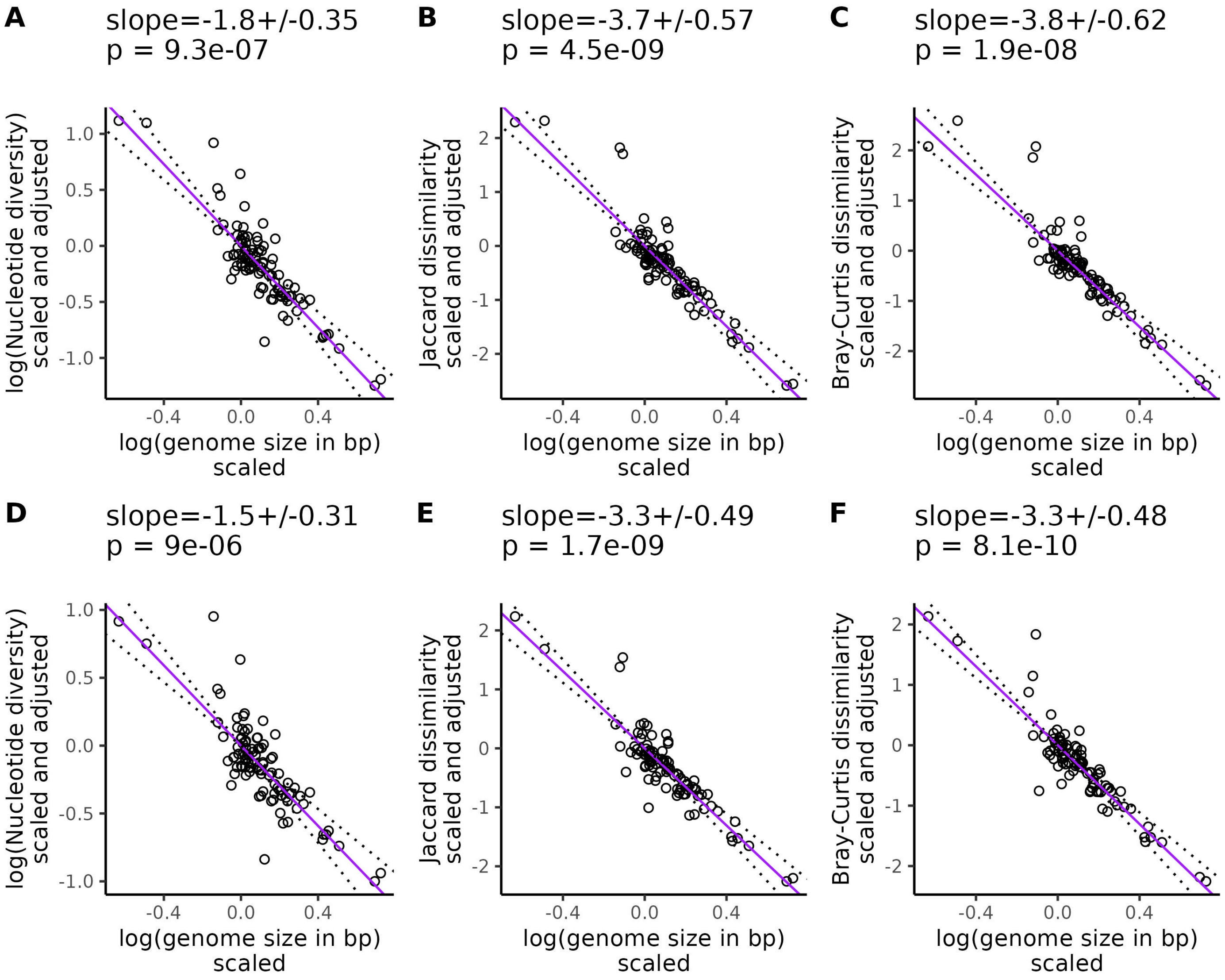


Figure S24: **Partial phylogenetic regression between diversity and genome size, controlling for WCVP range size.** Each point is a species and only species with > 0.5x mean coverage and > 1000 variant sites were included in the regression. The regressions are organized according to whether invaded ranges were excluded (A-C) or included (D-F) in the range size estimates. Lines give the relationship between the pairs of plotted variables after controlling for mating system, life cycle habit, cultivation status, range size, and evolutionary history. The statistics across the top of each plot give the value of the slope of the lines ($\pm$ the standard error) and the p-value testing whether the slope differs from zero. Before fitting the line, each response variable was scaled to a standard normal distribution (mean = 0, variance = 1), then multiplied by the inverse of the Cholesky decomposition of the phylogenetic variance-covariance matrix to correct for phylogenetic relationships. Dotted lines show the partial regression slope $\pm$ one standard error. All logarithms are base 10.

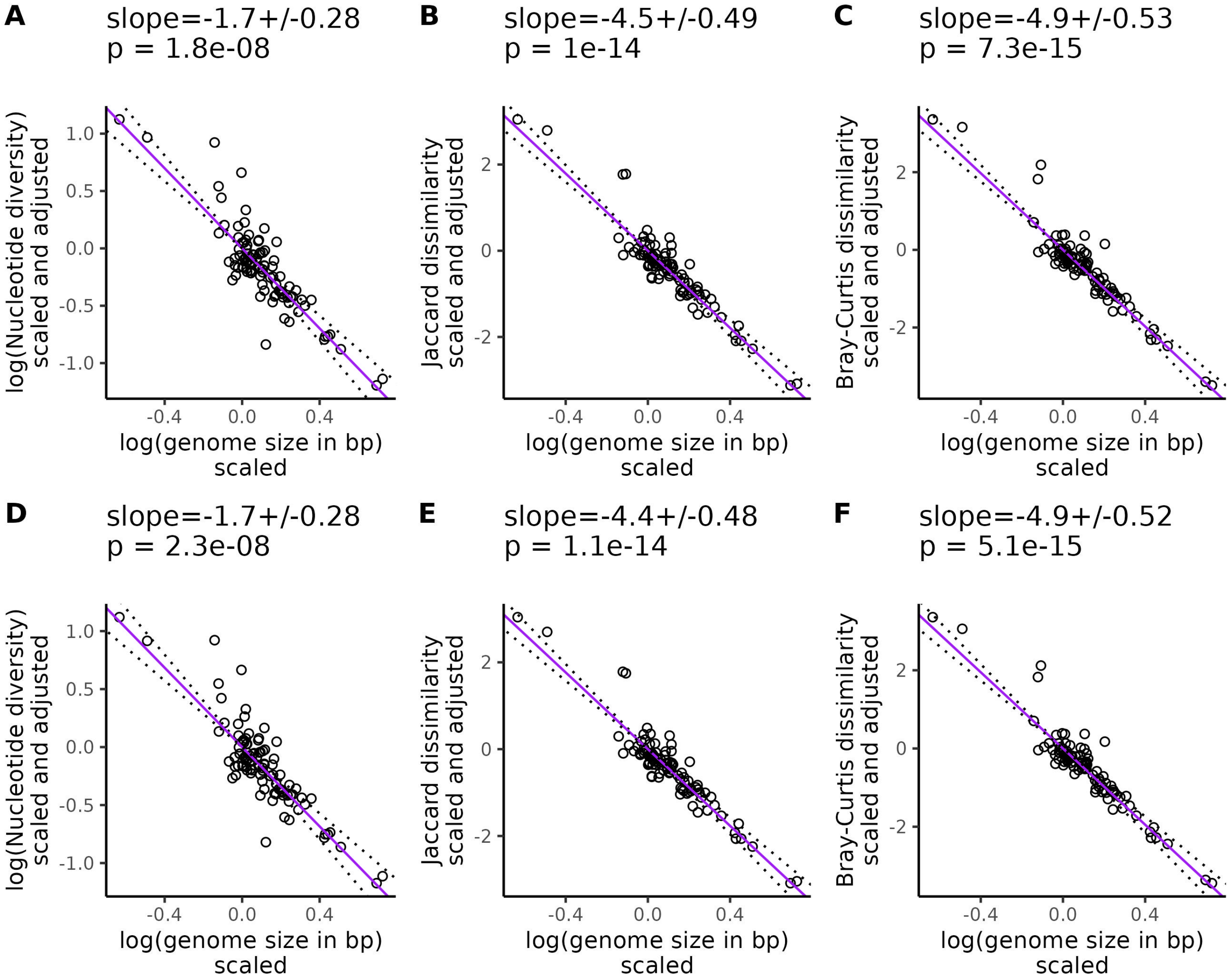


Figure S25: **Partial phylogenetic regression between diversity and genome size, controlling for GBIF range size.** Each point is a species and only species with $>$ 0.5x mean coverage and $>$ 1000 variant sites were included in the regression. The regressions are organized according to whether invaded ranges were excluded (A-C) or included (D-F) in the range size estimates. Lines give the relationship between the pairs of plotted variables after controlling for mating system, life cycle habit, cultivation status, range size, and evolutionary history. The statistics across the top of each plot give the value of the slope of the lines ($\pm$ the standard error) and the p-value testing whether the slope differs from zero. Before fitting the line, each response variable was scaled to a standard normal distribution (mean = 0, variance = 1), then multiplied by the inverse of the Cholesky decomposition of the phylogenetic variance-covariance matrix to correct for phylogenetic relationships. Dotted lines show the partial regression slope $\pm$ one standard error. All logarithms are base 10.

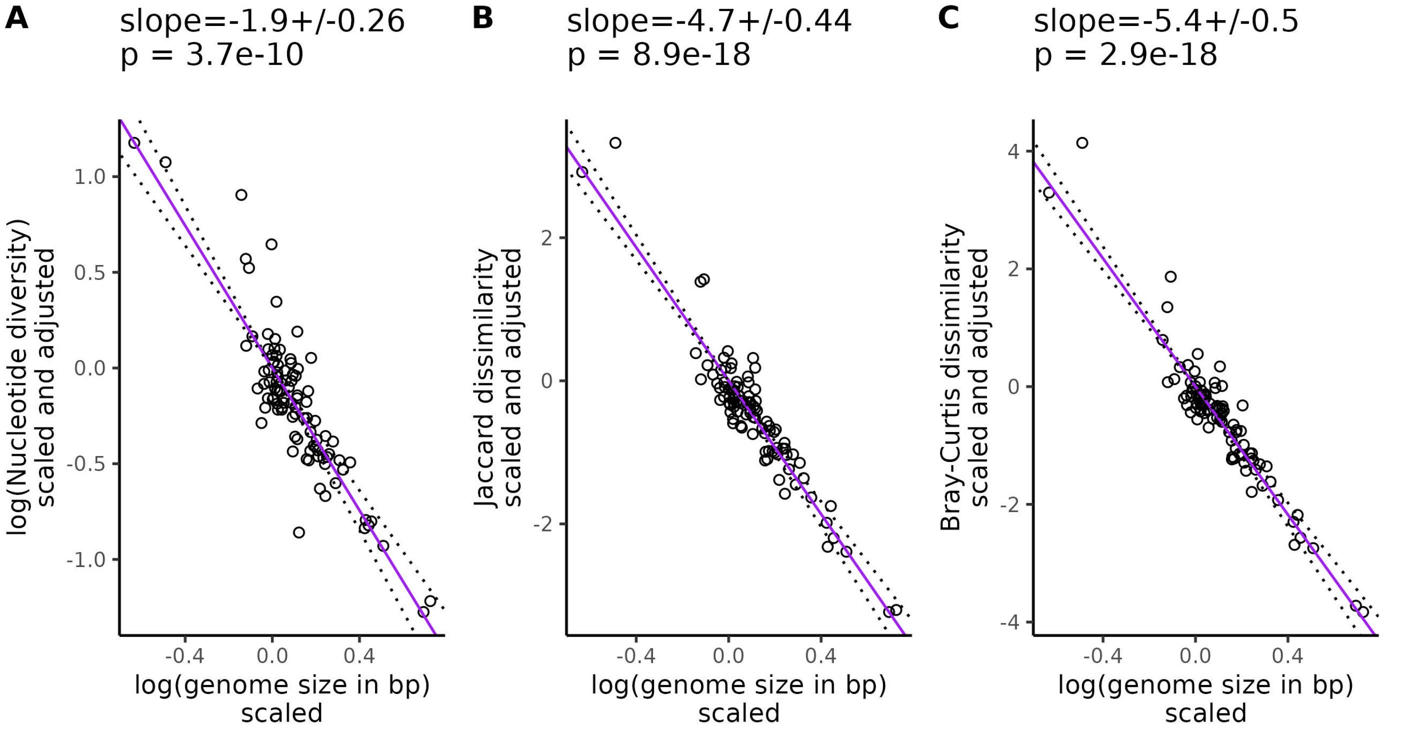


Figure S26: **Partial phylogenetic regression between diversity and genome size, controlling for height.** Each point is a species and only species with > 0.5x mean coverage and > 1000 variant sites were included in the regression. The regressions are organized according to whether invaded ranges were excluded (A-C) or included (D-F) in the range size-squared height ratio. Lines give the relationship between the pairs of plotted variables after controlling for mating system, life cycle habit, cultivation status, height, and evolutionary history. The statistics across the top of each plot give the value of the slope of the lines ($\pm$ the standard error) and the p-value testing whether the slope differs from zero. Before fitting the line, each response variable was scaled to a standard normal distribution (mean = 0, variance = 1), then multiplied by the inverse of the Cholesky decomposition of the phylogenetic variance-covariance matrix to correct for phylogenetic relationships. Dotted lines show the partial regression slope $\pm$ one standard error. All logarithms are base 10.
